# Telomere length and chromosomal instability for predicting individual radiosensitivity and risk via machine learning

**DOI:** 10.1101/2020.03.27.009043

**Authors:** Jared J. Luxton, Miles J. McKenna, Aidan M. Lewis, Lynn E. Taylor, Sameer G. Jhavar, Gregory P. Swanson, Susan M. Bailey

## Abstract

The ability to predict a cancer patient’s response to radiotherapy and risk of developing adverse late health effects would greatly improve personalized treatment regimens and individual outcomes. Telomeres represent a compelling biomarker of individual radiosensitivity and risk, as exposure can result in dysfunctional telomere pathologies that coincidentally overlap with many radiation-induced late effects, ranging from degenerative conditions like fibrosis and cardiovascular disease to proliferative pathologies like cancer. Here, telomere length was longitudinally assessed in a cohort of fifteen prostate cancer patients undergoing Intensity Modulated Radiation Therapy (IMRT) utilizing Telomere Fluorescence *in situ* Hybridization (Telo-FISH). To evaluate genome instability and enhance predictions for individual patient risk of secondary malignancy, chromosome aberrations were also assessed utilizing directional Genomic Hybridization (dGH) for high-resolution inversion detection. We present the first implementation of individual telomere length data in a machine learning model, XGBoost, trained on pre-radiotherapy (baseline) and *in vitro* exposed (4 Gy γ-rays) telomere length measures, to predict post-radiotherapy telomeric outcomes, which together with chromosomal instability provide insight into individual radiosensitivity and risk for radiation-induced late effects.

## Introduction

Radiation late effects are a broad class of negative and often permanent health effects experienced by cancer patients long after radiation therapy^1,2^, which can include cardiovascular disease^3^, pulmonary and arterial fibrosis^4^, cognitive deficits^5^, bone fractures^6^, and secondary cancers^7^. Such late effects are of particular concern for pediatric patients^8^, and risks for radiation late effects are highly dependent on patient-intrinsic factors as well, including genetics, age, sex, and lifestyle^1,2,9^. Therefore, identifying a patient’s specific risks for radiation late effects prior to radiotherapy is important for improving individual treatment planning and overall patient outcomes. A number of strategies for predicting risks for radiation late effects have been employed, which tend to irradiate patient-derived samples *in vitro* for monitoring of biomarker(s) to infer *in vivo* cellular and normal tissue *in vivo* responses to exposure^10^; e.g., evaluation of γ-H2AX foci kinetics^11,12^, apoptosis in normal blood lymphocytes^13^, and chromosome aberration frequencies^14–16^. Additionally, Genome Wide Association Studies (GWAS)^17,18^, sequencing^19^, and imaging studies (i.e radiogenomics^20^) have revealed promising putative markers that show promise for predicting risks for late effects. However, accurately predicting an individual patient’s response to radiotherapy and associated risk of developing adverse late health effects remains challenging in terms of cost-effectiveness, throughput, and predictive power, therefore new approaches are needed.

Telomeres are protective features of chromosomal termini that guard against genome degradation and inappropriate activation of DNA damage responses (DDRs)^21,22^. It is well established that telomeres shorten with cell division, oxidative stress^23^, and aging^24^. Telomeres also shorten with a host of lifestyle factors (e.g., nutrition^25^, exercise^26^, stress^27^) and environmental exposures (e.g. air pollution^28^, UV^29^) as well. Telomere length is a highly heritable trait, as is telomere length regulation^30–33^, supportive of individual variation in telomeric response to specific stressors. Interestingly, short telomeres have been proposed as hallmarks of radiosensitivity^34^, and ionizing radiation (IR) exposure has been shown to evoke both shortening and length**e**ning of telomeres^35–40^. Short telomeres are biomarkers and even effectors for a range of aging-related pathologies^41^, including cardiovascular disease (CVD)^42^, pulmonary fibrosis^42^, and aplastic anemia^43^, degenerative conditions also regarded as radiation late effects^44–46^. On the other hand, longer telomeres are associated with increased cancer risk, particularly for leukemias^47^, a common cancer following IR-exposure^48^. Thus, patients with shorter telomeres after radiation therapy are more likely to develop short telomere (degenerative) pathologies, while patients with longer telomeres following radiotherapy are at higher risk for developing proliferative pathologies (cancer).

Given that telomere length is influenced by a variety of genetic factors^30–33^ and exposures including IR exposure^35–40^, we reasoned that a patient’s telomeric outcome post-radiation therapy, rather than their pre-treatment (baseline) measures, would be most informative for assessing individual risks for radiation late effects and long-term health consequences. Furthermore, since patient-derived pre-radiation therapy samples irradiated *in vitro* provide an informative proxy for individual patient radiosensitivity and response *in vivo*^49–51^, an effective means to accurately predict an individual patient’s telomeric outcome post-radiation therapy could be developed, thereby improving personalized treatment strategies and individual outcomes.

Chromosome aberrations (CAs) are well-established biomarkers of IR-exposure^52^, associated with virtually all cancers^53^, and highly informative indicators of risk for radiation late effects, in particular, secondary cancers^14–16^. Ionizing radiation is exceptional in its ability to induce prompt double strand breaks (DSBs)^54^, damage that obligates a cellular response to address and resolve. Chromosome rearrangements result from the misrepair of such damage, and so provide a quantitative measure of cellular capacity for DNA repair^52^. In general, IR-induced CAs negatively impact cell survival and genome stability, resulting in senescence, apoptosis, and cancer^52^, respectively. Notably, chromosomal inversions and deletions have previously been proposed as signatures of radiation-induced secondary cancers^55^. Cytogenetic analysis however, is both time and labor intensive, often requiring that hundreds or even thousands of cells be scored, limiting its clinical utility^56^. We speculated that inclusion of an additional type of CA, specifically inversions, now possible using the strand-specific cytogenomic methodology of directional Genome Hybridization (dGH)^57^, might serve to reduce the number of cells required, while also informing potential risks for secondary cancers.

Significant advancements have also been made in the application of machine learning (ML) to a variety of scenarios, including predictions related to acute radiation toxicity^58^, treatment planning^59^, and secondary cancer risk post radiation therapy^60^. Extreme Gradient Boosting (XGBoost) is a powerful ML model that uses a gradient boosted ensemble of decision trees to learn complex relationships (linear and nonlinear) within data^61^. XGBoost has many translational applications, such as predicting future gastric cancer risk^62^, lung cancer detection^63^, and radiation-related fibrosis^64^. One potentially limiting caveat to ML is the requirement for extraordinarily large amounts of data to create robust, generalizable models. Telomere Fluorescence *in situ* Hybridization (Telo-FISH) is a cell-by-cell imaging-based approach for measuring telomere length capable of generating sufficient volumes of data for developing ML models; average experiments generate 200,000 −1,000,000 individual telomere length measurements^65^. Interestingly and to date, individual telomere length measurements (Telo-FISH, Q-FISH, flow-FISH, etc.) have not been utilized in ML models for risk predictions, despite the informative nature of such an approach.

Here we provide a proof-of-principle demonstration utilizing longitudinal analysis of telomere length and chromosomal instability in fifteen (15) prostate cancer patients undergoing Intensity Modulated Radiation Therapy (IMRT). We present the first implementation of individual telomere length (Telo-FISH) data in a ML model - XGBoost - and evaluate its ability to predict post-IMRT telomeric outcomes using individual patient’s pre-IMRT (baseline) and *in vitro* irradiated telomere lengths. Overall, results provide insight into predicting individual radiosensitivity and risk for radiation-induced late effects.

## Results

### Longitudinal analyses of telomere length associated with radiation therapy

Blood was collected from 15 prostate cancer patients undergoing IMRT at baseline (pre-IMRT), immediately post-IMRT (conclusion of treatment regimen), and three months post-IMRT. Baseline blood samples were split, half serving as the non-irradiated control (0 Gy), and the other half irradiated *in vitro* (4 Gy, Cs^137^ γ-rays) as a proxy for individual radiation response. The lengths of thousands of individual telomeres (n = 50 cells/patient/time point) were measured on metaphase chromosomes (lymphocytes stimulated from whole blood) by Telo-FISH at all time points (1: pre-therapy non-irradiated; 2: *in vitro* irradiated (4 Gy); 3B: immediately post-IMRT; and 4 C: three months post-IMRT) **(Fig 1A)**. For the overall cohort, differences in mean telomere length (MTL) between samples approached, but did not reach statistical significance (p = 0.059, repeated measures ANOVA). Relative to the pre-IMRT non-irradiated samples, overall MTL modestly increased after 4 Gy *in vitro* irradiation, and showed an even greater increase immediately after completion of the IMRT regimen, suggesting that increased MTL is an overall response to radiation exposure in this cohort. At three months IMRT, MTL for the cohort approached pre-IMRT levels.

**Figure 1.**
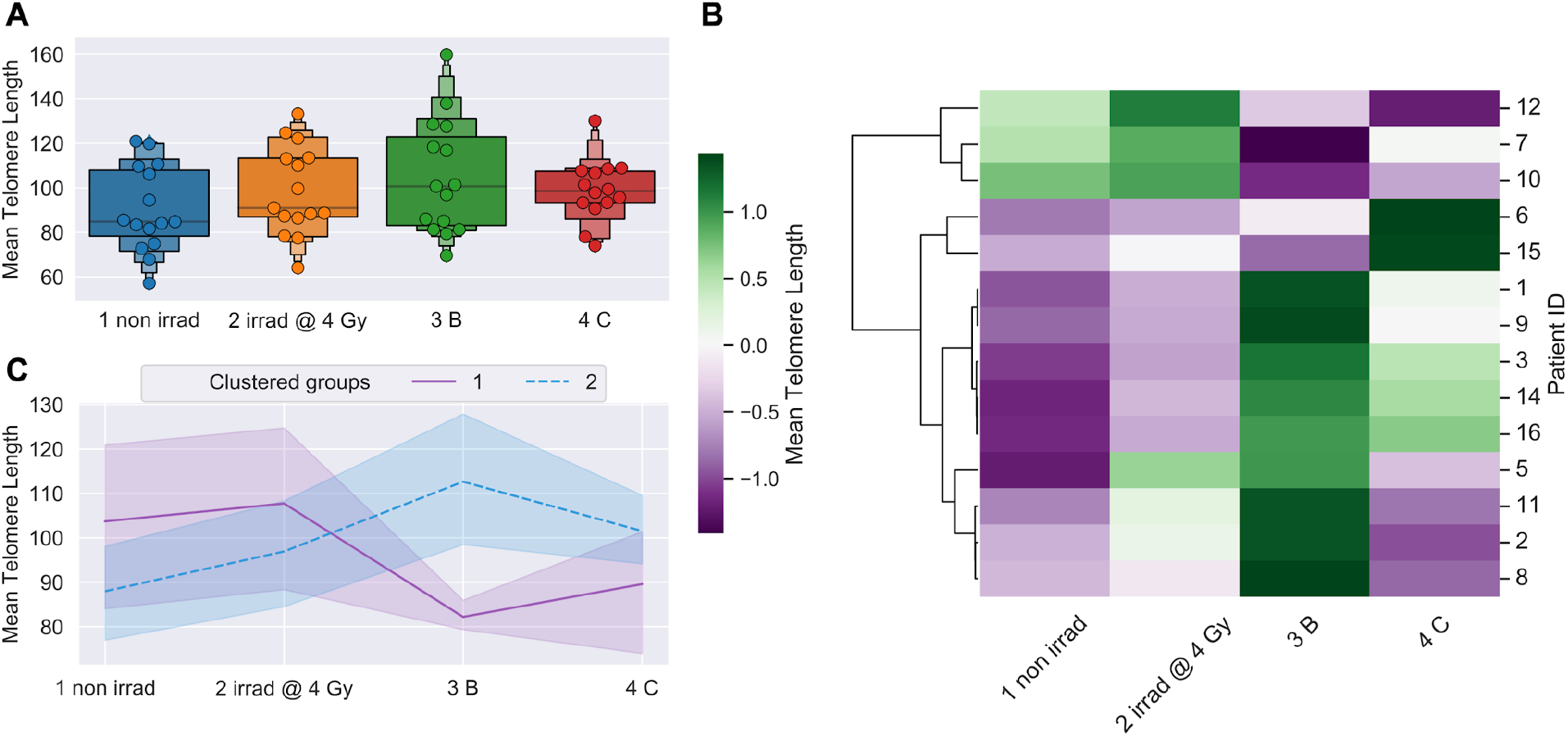
Telomere length dynamics (Telo-FISH). Mean telomere length expressed as relative fluorescence intensity. **A**) Time-course for all patients (n=15; 50 cells/patient/time point): 1 non irrad: pre-IMRT non-irradiated; 2 irrad @ 4 Gy: pre-IMRT *in vitro* irradiated; 3 B: immediate post-IMRT; and 4 C: 3 months post-IMRT. Boxes denote quantiles, horizontal grey lines denote medians. Telomere values were standardized using a pair of BJ1/BJ-hTERT control samples. **B**) Hierarchical clustering of patients by longitudinal changes in mean telomere length (z-score normalized). **C**) Time-course for clustered groups of patients (n=3, purple; n=11, blue); center lines denote medians, lighter bands denote confidence intervals. Patient ID 13 not clustered; 3 months post-IMRT sample failed to culture. Significance was assessed using a repeated measures ANOVA and post-hoc Tukey’s HSD test.

Complete blood counts (CBC) were also evaluated in the same samples, and longitudinal changes in patients’ MTL were negatively correlated (R^2^ = −0.126) with total peripheral white blood cell (WBC) counts (**Supp Fig 1A**). Longitudinal correlations between numbers of WBC types and MTL (all time points, for each patient) revealed a positive relationship with basophils (R^2^ = 0.278) and a negative relationship with lymphocytes (R^2^ = −0.294) (**Supp Fig 1B**). Furthermore, longitudinal correlations between MTL and the proportions of lymphocyte subgroups (all time points, for each patient) revealed positive relationships with natural killer (NK) and CD4 cells (R^2^ = 0.408, 0.282), and negative relationships with CD8 and CD19 cells (R^2^ = −0.251, −0.288) (**Supp Fig 1C**). These results support the notion that the overall changes in MTL associated with radiation exposure, specifically apparent telomere elongation, could be at least partially due to cell killing and shifts in lymphocyte populations, as previously proposed^36^.

### Telomere length dynamics revealed individual differences in radiation response

We hypothesized that groups of patients would cluster based on differential telomeric responses to radiation therapy, with sub-groups displaying either shorter or longer MTL post-IMRT. Clustering patients by longitudinal changes in MTL revealed two broad trends over time (**Fig 1B**). Patients that clustered in group 1 (n=3) had relatively longer MTL at baseline (pre-IMRT), and showed a dramatic, persistent *decrease* in MTL post-IMRT (**Fig 1C**). Those patients that clustered in group 2 (n=11) had relatively shorter MTL at baseline, and showed a dramatic, sustained *increase* in MTL post-IMRT (**Fig 1C**). Reduced MTL three months post-IMRT suggests increased risks for degenerative radiation late effects^42,43^, while increased MTL suggests increased risks for proliferative secondary cancers^47^.

In addition to MTL, Telo-FISH provides measures for many hundreds of individual telomeres, enabling generation of telomere length distributions and longitudinal analysis of shifts in populations of short and long telomeres^65^. For the overall cohort, numbers of short telomeres (yellow) decreased and numbers of long telomeres (red) dramatically increased three months post-IMRT (**Fig 2A**). When individual telomeres from patients in the MTL clustered group 1 (n=3) were combined, dramatic and persistent increases in the numbers of short telomeres post-IMRT were observed (**Fig 2B**), while MTL clustered group 2 patients (n=11) showed dramatic and persistent increases in numbers of long telomeres post-IMRT (**Fig 2C**). Again, patients with increased numbers of short telomeres are presumed to have increased risks for degenerative radiation late effects^42,43^, while those with increased numbers of long telomeres are at increased risk of secondary cancers^47^. Numbers of short and long telomeres were feature engineered (see Materials and Methods) from each patient’s individual telomere length data for further analysis.

**Figure 2.**
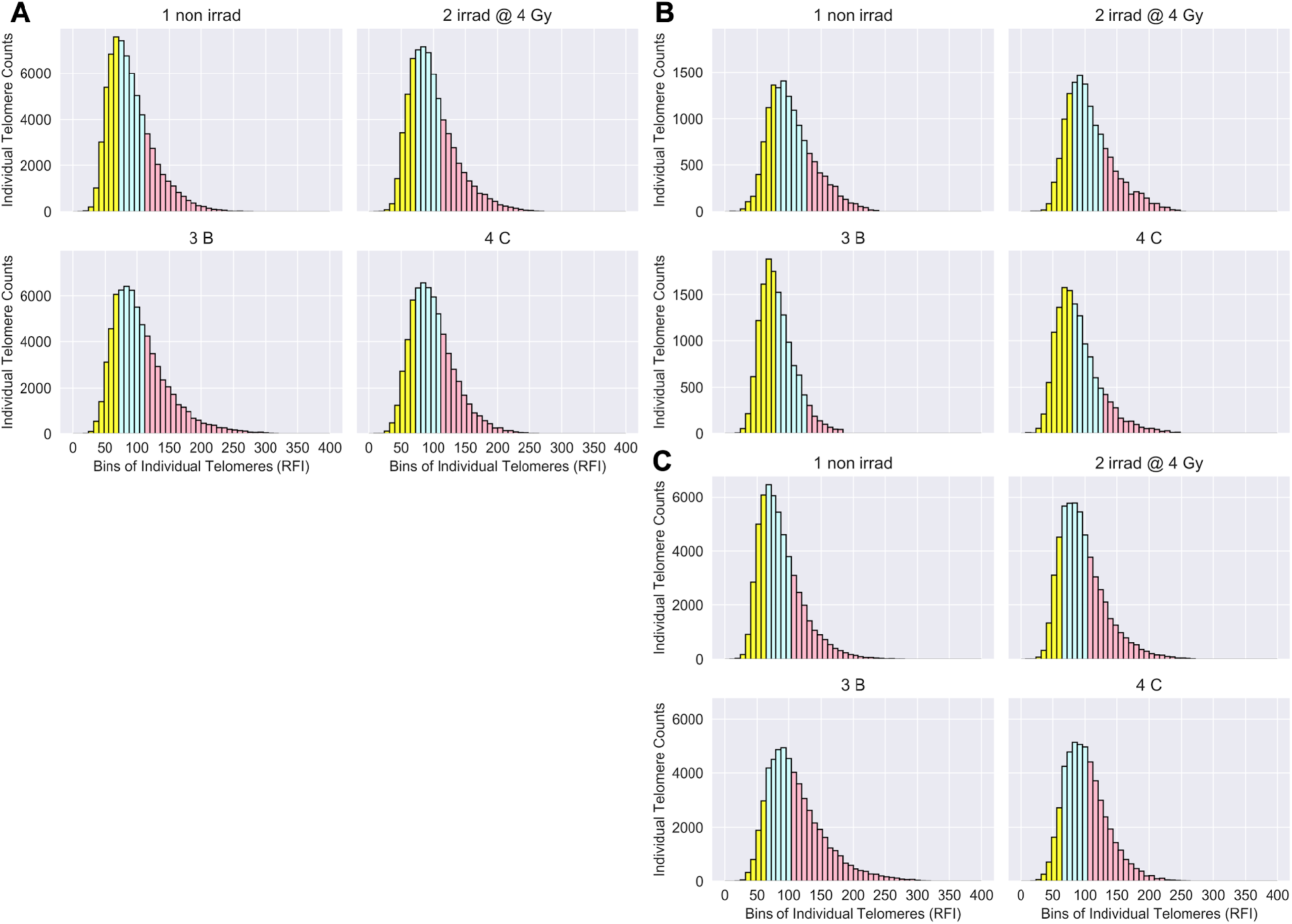
Telomere length distributions (Telo-FISH). Individual telomere length distributions of patients (n=15): 1 non irrad: pre-IMRT non-irradiated; 2 irrad @ 4 Gy: pre-IMRT *in vitro* irradiated; 3 B: immediate post-IMRT; 4 C: 3 months post-IMRT. RFI: Relative Fluorescence Intensity. Individual telomeres from the pre-therapy non-irradiated time point were split into quartiles, designating telomeres in bottom 25% in yellow, the middle 50% in blue, and top 25% in red. Quartile cut-off values, established by the distribution of the pre-therapy non-irradiated time point, were applied to subsequent time points to feature engineer the relative shortest (yellow), mid-length (blue), and longest (red) individual telomeres per time point. **A**) Individual telomeres for all patients (averaged) per time point. **B**) Individual telomeres for patients in mean telomere length clustered group 1 (n=3; aggregated) and **C**) group 2 (n=11; aggregated).

Differences in the average number of short and long telomeres between samples approached but did not reach statistical significance for the overall cohort (p<0.1; repeated measures ANOVA) (**Fig 3A**). We speculated that clustering patients by numbers of short or long telomeres would reveal longitudinal trends similar to those observed when clustering patients by MTL (**Fig 1B/C**). Clustering patients by longitudinal changes in numbers of short or long telomeres (**Fig 3B/D**) revealed two broad trends over time (**Fig 3C/E**). Clustered group 1 (n=3) showed a dramatic, sustained increase in numbers of short telomeres post-IMRT, with a corresponding decrease in numbers of long telomeres (**Fig 3C/E**). Clustered group 2 (n=11) showed a dramatic, nearly uniform decrease in numbers of short telomeres post-IMRT, with a corresponding increase in long telomeres (**Fig 3C/E**). Importantly, clustering patients either by MTL or by numbers of short or long telomeres post-IMRT identified the same three patients with shorter telomeres, and eleven with longer telomeres (**Fig 1B**, **Fig 3B/D)**.

**Figure 3.**
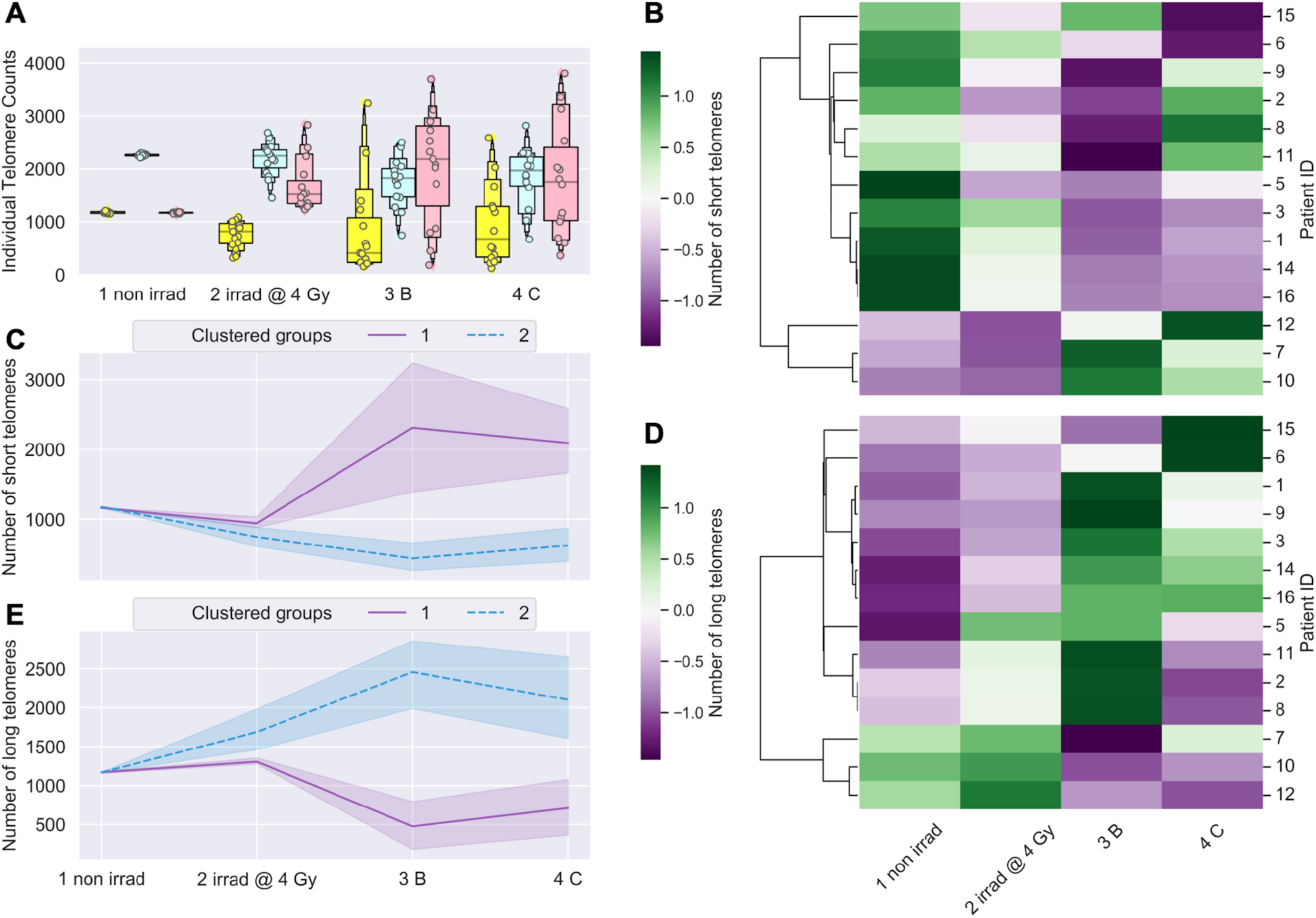
Longitudinal shifts in numbers of short and long telomeres (Telo-FISH). Numbers of short and long telomeres feature engineered from individual telomere length distributions per patient. 1 non irrad: pre-IMRT non-irradiated; 2 irrad @ 4 Gy: pre-IMRT *in vitro* irradiated; 3 B: immediate post-IMRT; 4 C: 3 months post-IMRT. Shortest (yellow), mid-length (blue), and longest (red) telomeres were feature engineered per patient (n=15) (see Materials and Methods). **A**) Counts of short, medium, and long telomeres, 4600 individual telomeres per patient (n=15) per time point. Significance was assessed for counts of short and long telomere using a squareroot transformation and a repeated measures ANOVA with post-hoc Tukey’s HSD test. Hierarchical clustering of patients by longitudinal changes in numbers of short telomeres **B**) and long telomeres **D**) (z-score normalized). Time-courses of patient groups (n=3, purple; n=11, blue) clustered by numbers of short **C**) and long **E**) telomeres; center lines denote medians and lighter bands denote confidence intervals. Patient ID 13 not clustered; 3 months post-IMRT sample failed to culture.

### Linear regression poorly predicted post-IMRT telomeric outcomes

Based on the two distinct groups identified by MTL and numbers of short and long telomeres three months post-IMRT (**Fig 1/3**), we hypothesized that pre-IMRT measurements of MTL and numbers of short and long telomeres could predict their respective post-IMRT outcomes using linear regression. For MTL, two linear regression models were created. The first used only MTL from pre-IMRT (baseline) non-irradiated samples as the independent variable, and the second used MTL from both the non-irradiated and *in vitro* irradiated pre-IMRT samples as independent variables for predicting post-IMRT MTL (**Fig 4A**). The R^2^ values for the two models were 0.161 and 0.165 respectively (**Fig 4A**), evidence that linear regression poorly captured the relationship between pre- and post-IMRT MTL. For numbers of short and long telomeres, two linear regression models were similarly created. The models for short telomeres yielded R^2^ values of 0.433 and 0.554, and the models for long telomeres yielded R^2^ values of 0.046 and 0.208 (**Fig 4B/C**). While the models for numbers of short telomeres had modestly higher R^2^ values than those for MTL or long telomeres, all linear regression models performed too poorly to confidently predict telomeric outcomes.

**Figure 4.**
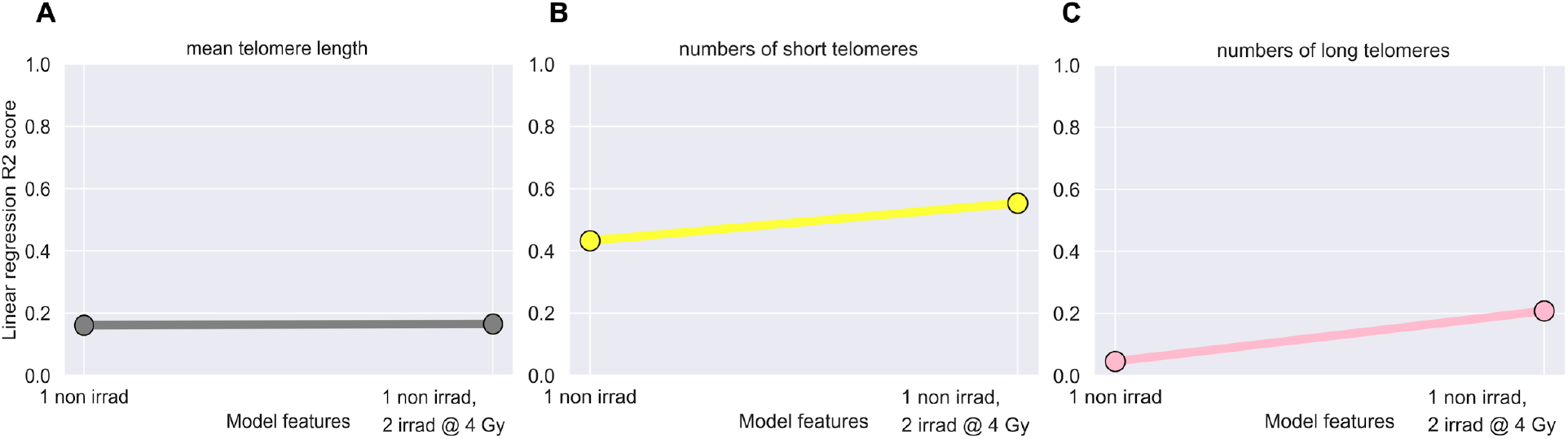
Linear regression models failed to predict post-IMRT telomeric outcomes. Ordinary least squares linear regression models were made using pre-IMRT telomeric data (Telo-FISH) from only the non-irradiated (1 non irrad) or also the *in vitro* irradiated (2 irrad @ 4 Gy) samples to predict 3 months post-IMRT telomeric outcomes. R2 values indicate the amount of variance in 3 months post-IMRT telomeric outcomes explained by the pre-therapy sample data. Models made using mean telomere length **A**) and numbers of short **B**) and long **C**) telomeres.

### Development of XGBoost machine learning models for accurate prediction of post-IMRT telomeric outcomes

The fact that linear regression poorly predicted post-IMRT telomeric outcomes could be due to the low number of observations (n=14), and/or the nonlinearity of telomere length dynamics (changes over time) in response to radiation exposure (**Fig 1-4).** We sought an alternative approach that could effectively utilize our vast dataset of pre-IMRT individual telomere length measurements (n=128,800), and also capture the nonlinearity of telomeric responses. Considering that XGBoost had recently been used to predict cancer risk and radiation-induced fibrosis using patient data^61–64^, we hypothesized that XGBoost models could be trained with pre-IMRT individual telomere length measurements to accurately predict post-IMRT telomeric outcomes.

Pre-IMRT (baseline) telomere length data required extensive preprocessing prior to training the XGBoost model for predicting three-month post-IMRT MTL (**Fig 5**). Data was reshaped into a matrix consisting of 128,800 rows (one for each individual telomere measurement) and four columns: patient ID, individual telomere length value, label denoting pre-IMRT sample of origin (non-irradiated or *in vitro* irradiated), and three-month post-IMRT MTL (**Supp Table 1A**). Reshaped data was randomly shuffled and stratified by patient ID and sample of origin, then split into training (80% of total) and test (20% of total) data sets. Shuffling guarded against order of measurement bias (Telo-FISH image acquisition), while stratifying ensured equivalent numbers of individual telomeres from each patients’ pre-IMRT samples (nonirradiated vs. *in vitro* irradiated) in the training and test data sets. Patient IDs were stripped from the training and test data sets, and individual telomeres from the non-irradiated and *in vitro* irradiated samples were encoded as 0 and 1 to denote sample origin (**Supp Table 1B**). XGBoost model hyperparameters were optimized using a randomized hyperparameter search^66^.

**Figure 5.**
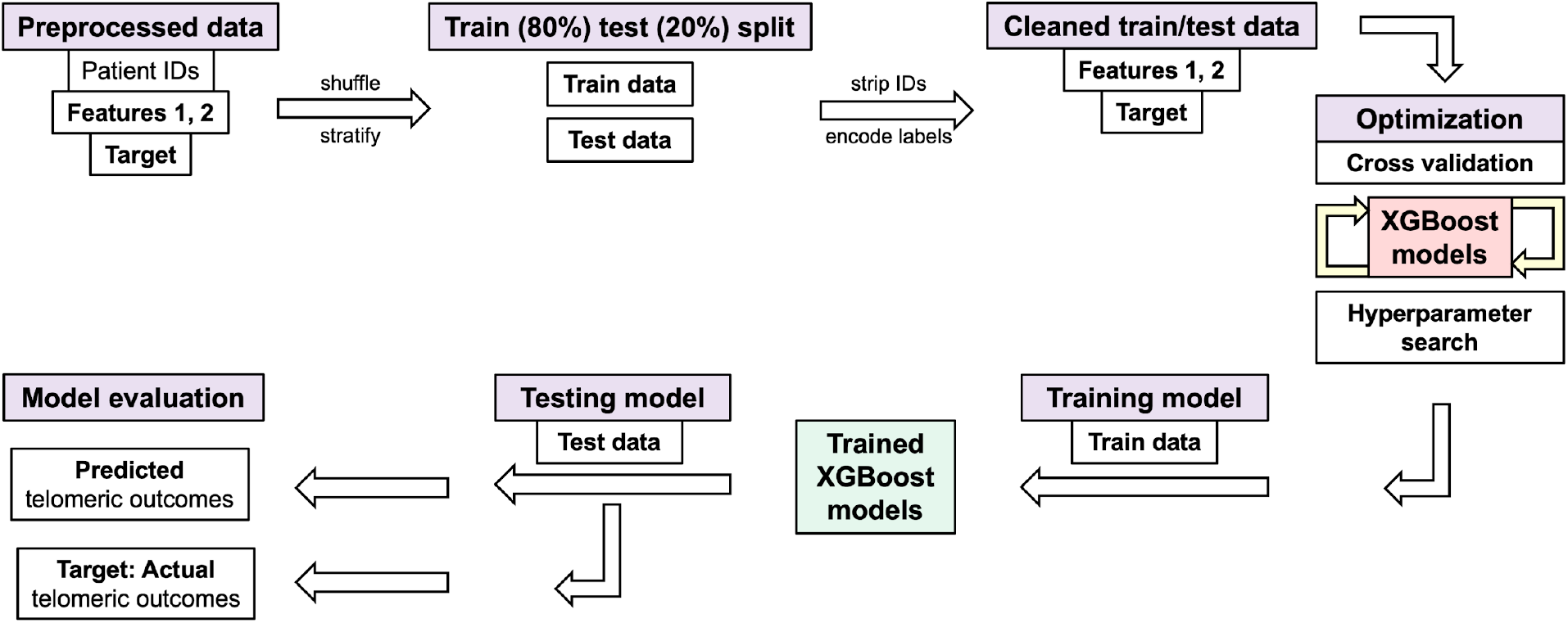
Processing of Telo-FISH data for training and testing XGBoost models. Schematic for our machine learning pipeline with individual telomere length data (Telo-FISH). Preprocessed data: Feature 1: pre-IMRT individual telomere lengths measurements (n=128,800); Feature 2: pre-IMRT sample labels (non-irradiated, *in vitro* irradiated, encoded as 0/1); Target: 3 months post-IMRT telomeric outcomes (mean telomere length or numbers of short and long telomeres). Data is randomly shuffled and stratified (by patient ID and pre-therapy sample origin) and split into training (80%) and testing (20%) datasets; patient IDs are stripped after splitting. Five-fold cross validation was used, and models were evaluated with Mean Absolute Error (MAE) and R2 scores between predicted and true values in the test set. See Materials and Methods and Code availability for model hyperparameters and implementations in Python.

XGBoost model performance was evaluated across the training data set using five-fold cross validation^67^. Mean absolute error (MAE), the mean of all differences between predicted and actual values of mean telomere length, was used to assess the model’s performance and ability to generalize to new data (**Supp Table 2A**). Five-fold cross validation on the full training data set yielded an average MAE of 3.233 with a standard deviation of 0.052 (**Supp Table 2A**), suggesting that the model was not overfitting to portions (folds) of the training data and that it could generalize to new data. Model performance was also evaluated when training across variable numbers of individual telomere measurements (n=100 to 103,040) (**Supp Table 2A**). After training the XGBoost model on the full training data set, the model was challenged to predict three-month post-IMRT MTL using new data - the test data set. The XGBoost model predictions for MTLs in the test set matched the true values with an R^2^ value of 0.882 (**Supp Table 2A**). Averaging predictions per patient for three-month post-IMRT MTL in the test set increased the R^2^ value to 0.931 (**Fig 6A**). Together, these results demonstrate that the XGBoost model learned the nonlinear relationships between pre-IMRT individual telomere length data and three-month post-IMRT MTLs (training data set), and generalized to new data (test data set) with highly accurate predictions.

**Figure 6.**
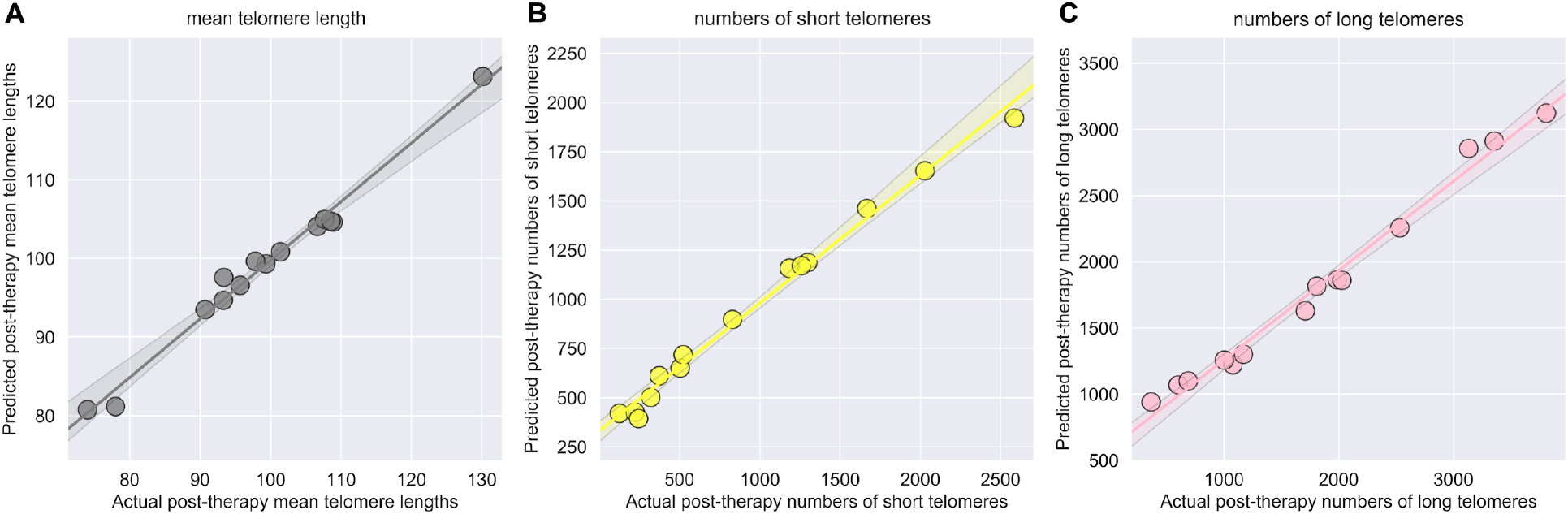
High performance of XGBoost models for predicting post-IMRT telomeric outcomes. XGBoost models were trained on pre-IMRT individual telomere length measurements (n=103,040, Telo-FISH) to predict 3 months post-IMRT telomeric outcomes. Trained XGBoost models were challenged with the test set (new data, n=25,760 individual telomeres) to predict 3 month post-IMRT telomeric outcomes. XGBoost predictions were averaged on a per patient basis for mean telomere length **A**) and numbers of short **B**) and long **C**) telomeres. R2 values between averaged predictions and actual values were 0.931 (A), 0.877 (B), and 0.890 (C).

Pre-IMRT individual telomere length data was also processed and reshaped for training separate XGBoost models to predict numbers of short or long telomeres three months post-IMRT (**Fig 5**, **Supp Table 1C/E**). Reshaped data was split into training (80%) and test (20%) data sets and shuffled and stratified in an identical manner as described for MTL (**Supp Table 1D/F**). Hyperparameters of the XGBoost models were optimized using a randomized search^66^, and the model’s performance and generalizability were analyzed using five-fold cross validation^67^ with a MAE error metric. For XGBoost models for short telomeres, five-fold cross validation on the full training data set yielded an average MAE of 232.3 with a standard deviation of 5.870 (**Supp Table 2B**), while XGBoost models for long telomeres yielded an average MAE of 326.0 and standard deviation of 3.93, suggesting that both models were reasonably good at fitting the data and likely to generalize to new data (**Supp Table 2C**). Model performance was also evaluated using variable numbers of training data (n=100 to 103,040). Fully trained XGBoost models were challenged with predicting three-month post-IMRT numbers of short or long telomeres in the test set, and predictions matched the true values with an R^2^ value of 0.814 and 0.827, respectively (**Supp Table 2B/C**). Averaging predictions per patient for post-IMRT numbers of short or long telomeres increased the R^2^ value to 0.877 and 0.890, respectively (**Fig 6B/C**). These results suggested that the XGBoost models learned the relationships between pre-IMRT individual telomere length data and three-month post-IMRT numbers of short or long telomeres (training data set), and effectively generalized to new data (test data set).

### Longitudinal analyses of chromosomal instability associated with radiation therapy

Directional Genomic Hybridization (dGH) is a cytogenomics, fluorescence-based methodology for high-resolution detection of chromosome aberrations (CAs) missed even by sequencing^68^, particularly inversions^57,69^. We hypothesized that the increased efficiency of dGH for detecting inversions would facilitate scoring fewer metaphase spreads (n=30/time point/patient) than traditional cytogenetic techniques^56^, while still retaining superb sensitivity to individual chromosomal instability, and thus the ability to infer patients at higher risks for secondary cancers. Many significant differences in frequencies of IR-induced rearrangements were observed (**Fig 7A-D**), with inversions occurring at the highest frequencies, consistent with expectations^57,69^. Interestingly, overall average frequencies of inversions at three months post-IMRT were comparable to the *in vitro* irradiated samples (**Fig 7A**). Frequencies of translocations, dicentrics, and chromosome fragments (deletions) were highest after *in vitro* irradiation, and remained relatively high immediately post-IMRT (**Fig 7B-D)**. High frequencies of translocations and chromosome fragments persisted post-IMRT (**Fig 7B/D**), while dicentrics decreased somewhat (**Fig 7C**). Frequencies of sister chromatid exchanges (SCE) did not significantly change over time, consistent with expectation and exposure. Notably, elevated frequencies of CAs at three months post-IMRT suggested ongoing genomic instability in the overall cohort^52,53,55^ (**Fig 7A-D**).

**Figure 7.**
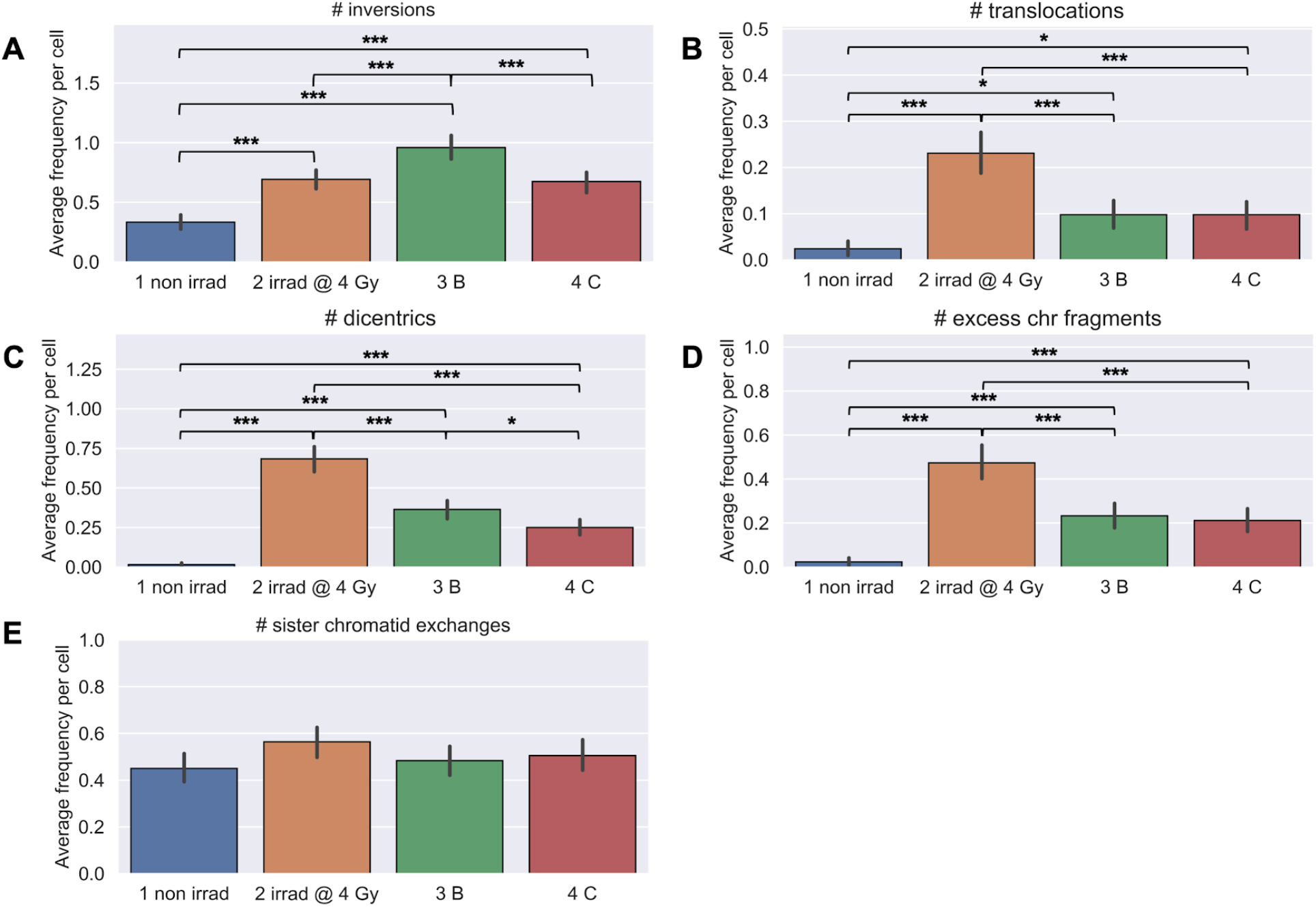
Longitudinal analyses of chromosomal instability (dGH). Chromosome aberrations were scored using directional Genomic Hybridization (dGH) in cultured T-cells harvested in metaphase (n=30/patient/timepoint) from whole blood of patients (n=15). 1 non irrad: pre-IMRT non-irradiated; 2 irrad @ 4 Gy: pre-IMRT *in vitro* irradiated; 3 B: immediate post-IMRT; 4 C: 3 months post-IMRT. Counts of inversions and translocations (A/B) were adjusted for clonality, where identical aberrations between cells are noted but scored only once. Excess chr fragments: counts of chromosome fragments per cell after subtracting 1 count per n observed dicentrics. **A**) inversions, **B**), translocations, **C**) dicentrics, **D**) chromosome fragment, and **E**) sister chromatid exchanges. Significance was assessed for average aberration frequencies using a repeated measures ANOVA and post-hoc Tukey’s HSD test. p<0.05, p<0.01, p<.001 = *, **, ***.

Significant changes in frequencies of IR-induced rearrangements were also coincident with numbers of peripheral blood lymphocytes. Longitudinal correlations between patients’ average frequencies of CAs and numbers of peripheral blood lymphocytes (all time points) revealed strongly negative correlations (**Supp Fig 2A-D**). Frequencies of inversions and dicentrics had the highest negative correlations (R^2^ = −0.752, −0.751), indicating they were highly informative - and similar - markers for cell death. These results suggest that patients demonstrating chromosomal instability (specifically, elevated frequencies of inversions or dicentrics), also experience higher levels of cell killing (i.e., greater radiosensitivity) consistent with previous reports^70,71^.

Next, we hypothesized that clustering patients by longitudinal changes in CA frequencies (all samples) would reveal groups of patients with lower or higher frequencies of CAs, which would be indicative of individual chromosomal instability and radiosensitivity. When clustering patients by CA type, we observed groups of patients with differential responses only for inversions and chromosome fragments (deletions), which displayed increased frequencies immediately post-IMRT, suggesting increased chromosomal instability (**Fig 8A/D**, **Supp Fig 3A/B**). We note that the two patients with the highest post-IMRT frequencies of inversions (ID #16) and chromosome fragments (ID #6), also had very high post-IMRT MTLs; both biomarkers suggestive of increased risks for secondary cancers^14–16,47^ **(Fig 1C, Fig 8A/D**, **Supp Fig 3A/B**).

**Figure 8.**
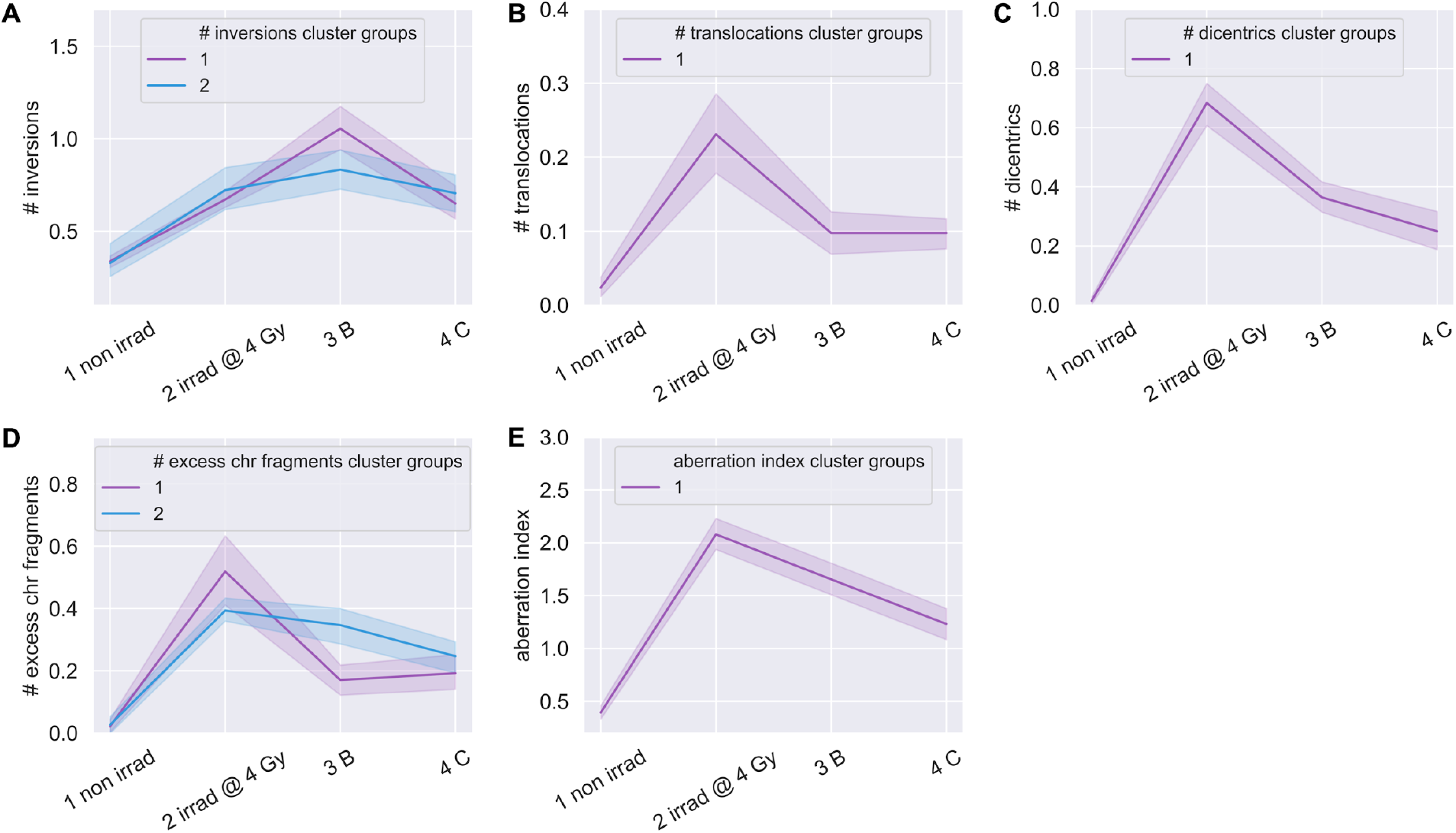
Clustering of patients by chromosome aberration frequencies. Time-courses for groups of patients hierarchically clustered into discrete groups (blue/purple) per aberration type. 1 non irrad: pre-IMRT non-irradiated; 2 irrad @ 4 Gy: pre-IMRT *in vitro* irradiated; 3 B: immediate post-IMRT; 4 C: 3 months post-IMRT. Excess chr fragments: counts of chromosome fragments per cell after subtracting 1 count per n observed dicentrics. Aberration index is created by summing all aberrations (A-D) per cell. Center lines denote medians and lighter bands denote confidence intervals. Clustered groups of patients for inversions **A**), translocations **B**), dicentrics **C**), chromosome fragments **D**), and aberration index **E**).

Other CA types had longitudinal responses that were relatively uniform between patients and did not cluster patients (**Fig 8B/C**, **Supp Fig 4A/B**). We hypothesized that while individual types of CAs failed to cluster patients into groups, individual patients may show lower or higher frequencies of CAs. To determine if some patients showed a general susceptibility to chromosomal instability, we feature engineered ‘aberration index’ by summing all types of CAs (less sister chromatid exchanges) (**Fig 8A-D**) per cell for all time points. As indicated by the aberration index, groups of patients with lower or higher total CA frequencies were not observed (**Fig 8E**, **Supp Fig 4C**). These results in conjunction with the telomere length data, identified two patients (ID #s 6, 16) at potentially increased risks for secondary cancers^14–16,47^, and are supportive of inversions and deletions being more informative than other CA types for predicting IR-induced secondary cancers, consistent with prior reports^55^. These results also indicate that the numbers of cells scored were too low (n=30) to detect significant differences in individual patient susceptibility to chromosomal instability in general.

### Linear regression poorly predicted radiation-induced chromosomal instability

We speculated that pre-IMRT CA frequencies could be predictive of post-IMRT frequencies. Two linear regression models were made for each CA type to predict post-IMRT frequencies; the first used only the pre-IMRT (baseline) non-irradiated sample CA frequency, and the second used CA frequencies from both pre-IMRT non-irradiated and *in vitro* irradiated samples. The models showed poor predictive power overall, and although inclusion of the *in vitro* irradiated sample data improved performance overall, both models were insufficient for predicting post-IMRT CA frequencies with confidence (**Fig 9A-E**). The model for dicentrics performed best, with an R^2^ score of 0.514 when using data from both irradiated and nonirradiated baseline samples. These results suggest that while *in vitro* irradiated sample data added predictive power, the number of cells scored per time point/patient (n=30) was too low to enable accurate predictions of individual patient outcomes regarding CAs frequencies post-IMRT using linear regression.

**Figure 9.**
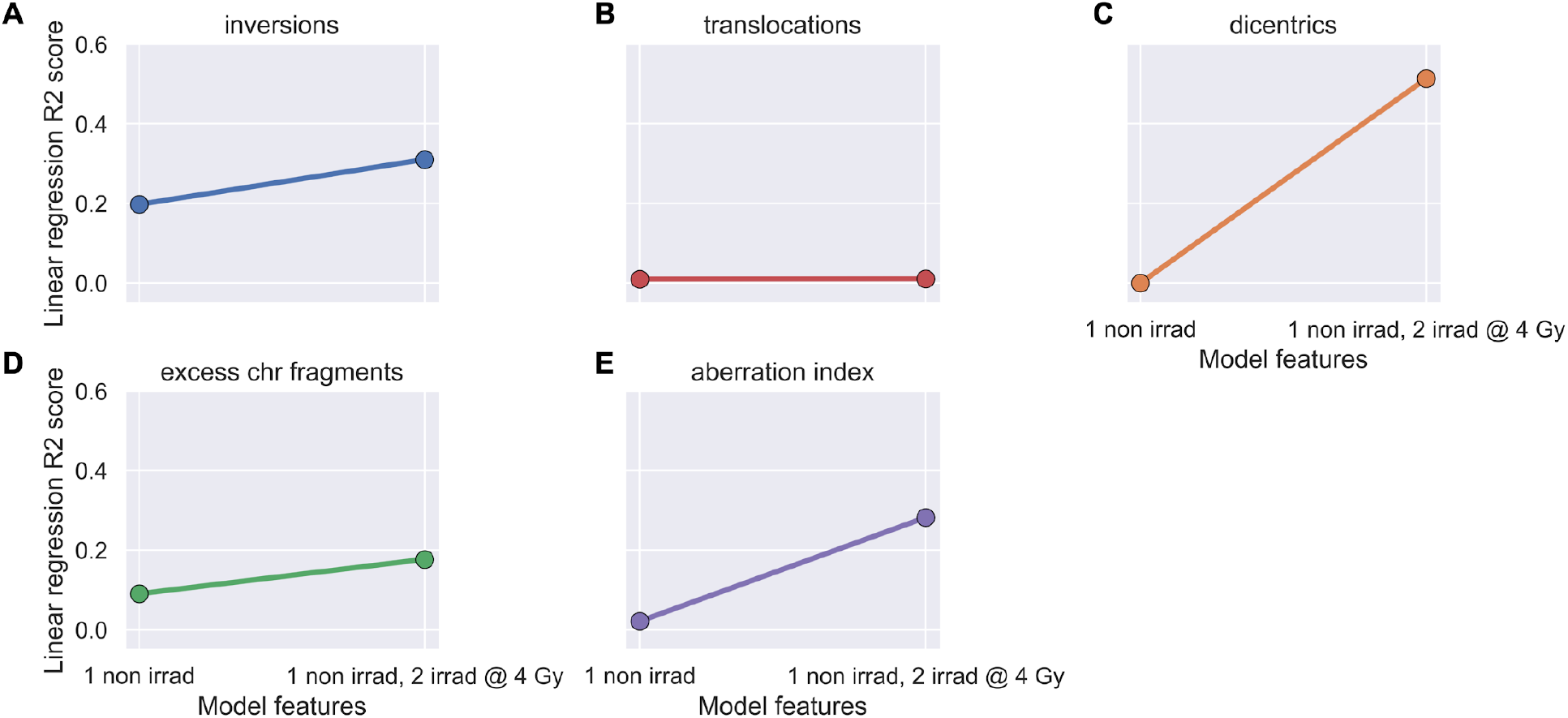
Linear regression models failed to predict post-IMRT chromosome aberration frequencies. Ordinary least squares linear regression models were made using pre-IMRT average aberration frequencies from only the non-irradiated (1 non irrad) or also the *in vitro* irradiated (2 irrad @ 4 Gy) samples to predict 3 months post-IMRT average aberration frequencies. Excess chr fragments: counts of chromosome fragments per cell after subtracting 1 count per n observed dicentrics. Aberration index is created by summing all aberrations (A-D) per cell. R2 values indicate the amount of variance in late post-IMRT outcomes explained by the pre-IMRT sample data. Models made with inversions **A**), translocations **B**), dicentrics **C**), chromosome fragments **D**), and aberration index **E**).

### XGBoost machine learning models poorly predicted radiation-induced chromosomal instability

We attempted training XGBoost models using pre-IMRT (baseline) CA data to predict post-IMRT CA frequencies. Rather than using CA data per patient, which would be insufficient for model training (n=15), we used pre-IMRT CA frequencies on a per cell basis (n=840) to predict three-month post-IMRT average CA frequencies. Pre-IMRT CA frequency data was extensively processed prior to XGBoost model training (**Supp Fig 5, Supp Table 3A-D**), in a nearly identical manner as described for pre-IMRT telomere length data. The key difference was that CA data was reshaped to train XGBoost models with pre-IMRT CA count data per cell (n=672 cells) in order to predict three-month post-IMRT average CA frequencies. Separate datasets and XGBoost models were created for each type of CA.

XGBoost models for each type of CA were evaluated across their respective training sets using five-fold cross validation^67^ with a MAE metric. The cross-validation metrics for all XGBoost models with CA data suggested a failure of the models to learn relationships between pre-IMRT CA count data per cell and three-month post-IMRT average CA frequencies (**Supp Table 4A**). Furthermore, dramatic fluctuations in model performance were noted when running multiple iterations of cross-validation, again suggesting that the models failed to learn the relationships between the pre- and post-IMRT CA frequencies (**Supp Table 4A-C**). We attempted to improve model performance with many types of feature engineering (e.g boolean features), numerical transformations, and adjustments to model hyperparameters, none of which yielded meaningful improvements in any combination (data not shown). Regardless of poor model performance in cross-validation, we challenged the XGBoost models to predict post-IMRT average CA frequencies using pre-IMRT CA count data per cell in the test set (n=168 cells). In XGBoost model predictions for three month post-IMRT CA frequencies in the test set, none of the predictions matched the true values, with an R^2^ above 0.1 (**Fig 10A-E, Supp Table 4A-C**). These results indicate that either the amount of data was insufficient for training XGBoost models (n=840 cells at pre-IMRT), or the strategy of predicting post-IMRT average CA frequencies using pre-IMRT CA count data per cell was inherently faulty.

**Figure 10.**
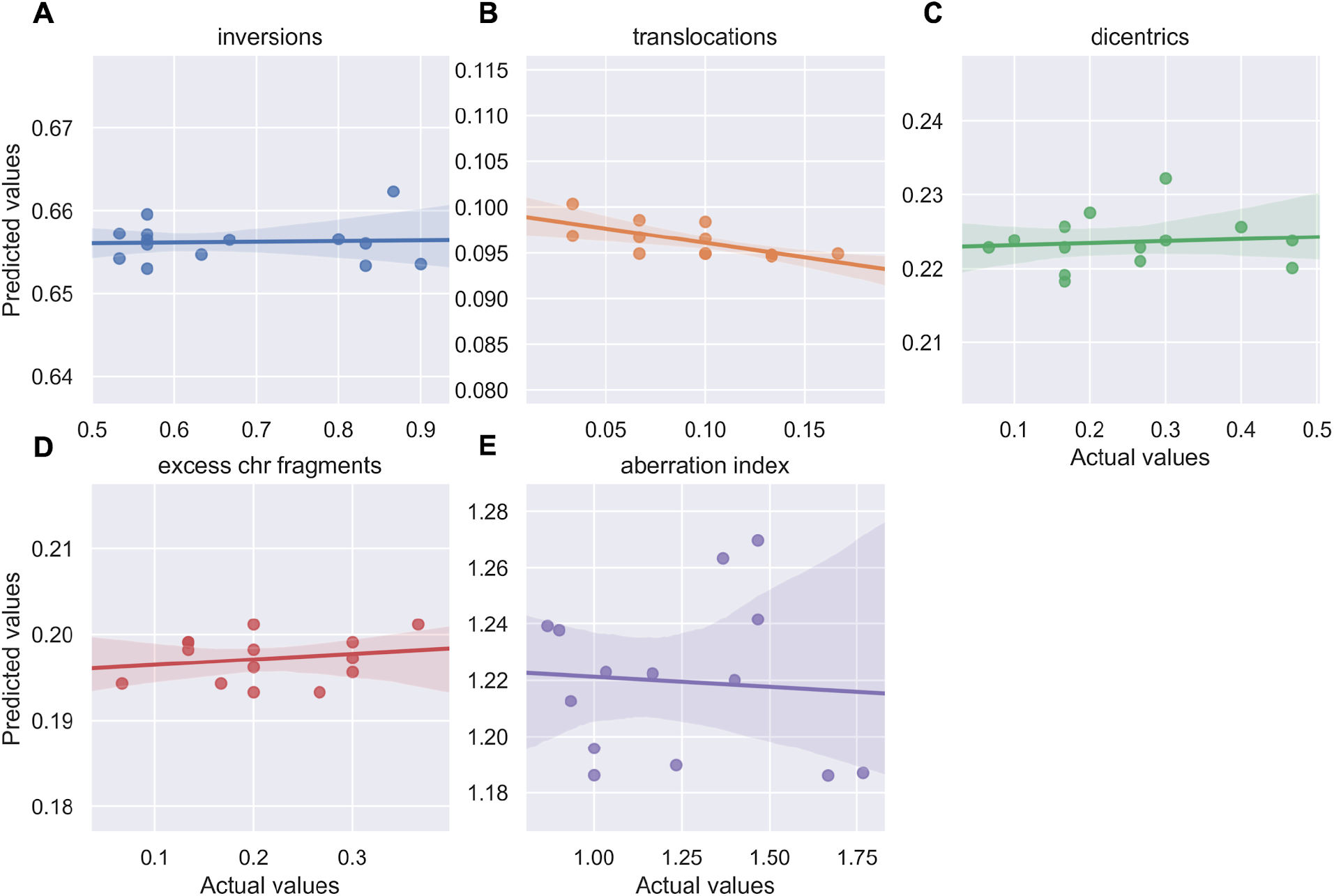
XGBoost models failed to predict post-IMRT chromosome aberration frequencies. XGBoost models were trained on pre-IMRT counts of different chromosome aberration types per cell (n=672) to predict 3 months post-IMRT average chromosome aberration frequencies. Trained XGBoost models were challenged with the test set (new data, n=168 cells) to predict 3 months post-IMRT average chromosome aberration frequencies. Excess chr fragments: counts of chromosome fragments per cell after subtracting 1 count per n observed dicentrics. Aberration index is created by summing all aberrations (A-D) per cell. XGBoost predictions were averaged on a per patient basis for inversions **A**), translocations **B**), dicentrics **C**), chromosome fragments **D**), and aberration index **E**). For all models, R2 values between averaged predictions and actual values did not exceed 0.100.

## Discussion

Better prediction of a cancer patient’s individual response to radiation therapy and risk for developing adverse late health effects remains a prime objective for the treatment modality in general^1–7^, and particularly in regard to pediatric patients^8^. Over recent years, a variety of approaches for predicting radiation late effects have been developed^10–20^, albeit with varying degrees of compromise between cost-effectiveness, throughput, and predictive power. One notable and extremely promising exception is the use of ML models, which can leverage extensive amounts of patient data to make accurate predictions of treatment outcomes^58–60,62–64^.

Predicting a patient’s telomeric response to radiation therapy is of clinical interest for predicting risks of radiation late effects, as shorter telomeres confer radiosensitivity^34^ and increase risk of degenerative late effects (CVD^42^, pulmonary fibrosis^42^, aplastic anemia^43^), while longer telomeres increase risk for secondary cancers, particularly leukemias^47^. Given that telomeric responses to radiation exposure can be highly dynamic^35–40^ and vary between individuals (**Fig 1-3**), a framework for predicting a patient’s particular telomeric responses to radiation therapy is critical for utilizing telomere length as a biomarker for radiation late effects. Here, we demonstrate the feasibility of using ML to accurately predict an individual patient’s telomeric response to radiation therapy. We successfully implemented individual telomere length data in a machine learning model, XGBoost, for highly accurate predictions of post-IMRT telomeric outcomes (**Fig 5/6, Supp Table 3**). The ML models and Telo-FISH methods used are fully available, providing a valuable resource for continued research into telomere length as a biomarker for radiation late effects associated with any manner of exposure.

The possibility of improving assessment of chromosomal instability and associated risk for development of secondary cancers following radiation therapy^14–16^ was also explored utilizing dGH, which facilitated inversion detection at higher resolution than traditional cytogenetic assays^57,69^. Indeed, inversions were observed at higher frequencies than other types of CAs both before and after radiation therapy (**Fig 7A**), consistent with prior reports^57,69^. Groups of patients with increased frequencies of chromosomal inversions and fragments (deletions), previously proposed signatures of radiation-induced cancers^55^, were also observed three months post-IMRT (**Fig 8A/D**). Two patients from these groups had very high MTLs three months post-IMRT as well, also supportive increased risks for secondary cancers^14–16,47^. We attempted to derive some predictive value from CA data with linear regression and XGBoost implementations, but both efforts were summarily unsuccessful; the low numbers of cells scored per patient likely subverted successful predictions from the data.

Although we were unable to predict post-IMRT changes in CA frequencies, our observations of strong correlations between patients’ average frequencies of CAs and changes in peripheral blood lymphocyte counts associated with IMRT indicated that our approach of detecting inversions for evaluating chromosomal instability was fundamentally correct. Patients with higher levels of radiation therapy-induced chromosomal instability also experienced increased levels of cell death; i.e., they exhibited individual radiosensitivity^70,71^. Inversions and dicentrics in particular had strong, negative correlations with lymphocytes cell counts (R^2^ = −0.752, −0.751) (**Supp Fig 2A/B**).

Relationships between peripheral blood cell count data and MTL were also observed. Counts of peripheral white blood cells (WBCs) were negatively correlated with MTL associated with IMRT (R^2^ = −0.126), supportive of shorter telomeres contributing to cell killing/radiosensitivity (**Supp Fig 1A**). When parsing WBCs by sub-type, a stronger negative relationship between MTL and lymphocyte counts was seen (R^2^ = −0.294). When parsing lymphocytes by sub-type and correlating MTL with the proportions of cell-types, we observed positive correlations with NK and CD4 cells (R^2^ = 0.408, 0.282), and negative correlations with CD8 and CD19 cells (R^2^ = −0.251, −0.288). These results support our previously proposed supposition that the observed changes in MTL associated with radiation exposure could be partially due to changes in peripheral blood lymphocyte cell populations^36^.

Longitudinal assessment of individual telomere length by Telo-FISH in cancer patients undergoing IMRT facilitated demonstration of XGBoost as the ML model of choice for predicting telomeric outcomes post-IMRT. Given the notion that risks for radiation late effects occur on a spectrum^1–8^, and the differential telomeric responses between individuals and radiation modalities, we posit that the true range of telomeric responses for radiation therapy patients in general is much broader than those observed here in this prostate cancer cohort (**Fig 1-3**). Thus, while our XGBoost models effectively generalized to new data within our experimental design (similar patient sex, radiation modality, cancer type, etc.) (**Fig 6**, **Supp Table 3**), it’s unlikely that our trained models, in their current iteration, would generalize to data collected under different experimental parameters. Moreover, with regard to measurement of individual telomere lengths for training XGBoost models, Telo-FISH could readily be interchanged with comparable assays (Q-FISH, flow-FISH), which may provide higher throughput. Additionally, the ML approaches described here were not strictly dependent upon XGBoost, and could be conducted using other machine learning models and frameworks (e.g., random forests, kNN). Our paradigm of training ML models with individual telomere length data for prediction of post-IMRT telomeric outcomes provides improved predictive power and novel insight into individual radiosensitivity and risk of late effects, as well as a general framework that could be deployed for radiation therapy patients regardless of cancer type, radiation modality, or individual patient sex or genetic susceptibilities.

## Data availability

Raw and processed individual telomere length (Telo-FISH) data files and chromosome aberration score sheets (dGH) are available for download at https://github.com/Jared-Luxton/. All data processing pipelines and code was written in Python and stored in Jupyter notebooks at https://github.com/Jared-Luxton/. The Jupyter notebooks can be run within a web browser and are available for download.

## Acknowledgements

Funding from the Colorado Office of Economic Development and International Trade (OEDIT) Advanced Industry (AI) Bioscience Proof of Concept (POC) award program, Colorado State University (CSU) Ventures, and KromaTiD, Inc. are gratefully acknowledged. Graduate student fellowhips awarded by CSU’s Program of Research and Scholarly Excellence (PRSE), and the Cancer Biology and Comparative Oncology (CB&CO) PRSE to support quantitative Cell and Molecular Biology (qCMB) studies, are also gratefully acknowledged.

## Author contributions

S.M.B, G.P.S, and M.J.M conceived the study. G.P.S. and S.G.J. secured collection of the samples. J.J.L., A.M.L., and L.E.T participated in sample processing. J.J.L. performed the data analysis. J.J.L. prepared the figures and tables. J.J.L. and S.M.B. wrote the manuscript. All authors exchanged ideas and contributed to editing the manuscript.

## Competing interests

S.M.B. is a cofounder and scientific advisory board member of KromaTiD, Inc.

## Materials and Methods

### Patient consent, IMRT therapy information

With informed consent as per the institutional review board, 16 consecutive patients that were receiving pelvis and prostate or prostate fossa radiation therapy were asked to participate. No patient had received androgen ablation or chemotherapy to avoid confounding factors. One patient was found to have metastatic disease after consent and was removed from further study. A total of 15 patients provided consent and blood was obtained at pre-IMRT (baseline), immediately post-IMRT (the last week) and 3 months post-IMRT. Blood was subject to complete blood counts, and telomere length and chromosome aberration analyses. Radiation consisted of 54 Gy to the pelvic lymphatics, with a total of 70 Gy (n=11) or 78 Gy (n=3) to the prostate fossa. One patient underwent brachytherapy boost.

### Sample collection and processing for Telo-FISH and dGH

Peripheral blood was drawn and shipped in 10 mL sodium heparin tubes (Becton, Dickinson, and Co #367874) under ambient conditions to Colorado State University and received within 24 hours of blood draw. All heparinized blood samples were cultured in T-25 tissue culture flasks, at 1 parts blood per 9 parts Gibco PB-Max Karyotyping Medium (ThermoFisher #12557013), with 5.0 mM 5-bromo-deoxyuridine (BrdU) and 1.0 mM 5-bromo-deoxycytidine (BrdC) added to the medium as previously described^57^. Pre-IMRT blood samples were split into two fractions (non-irradiated and *in vitro* irradiated) with identical culturing conditions as other time point samples, and one fraction was irradiated in a Cs137 irradiator *in vitro* at a dose rate of 2.5 Gy/min for a total dose of 4 Gy (γ-rays). 48 hours after stimulation, KaryoMax Colcemid (ThermoFisher #15210040) was added (0.1 μg per mL of medium) for four hours of incubation, then metaphase chromosome spreads were harvested with standard cytogenetic protocols^72^. Prior to Telo-FISH and dGH, slides with metaphase chromosome spreads were subject to CO-FISH for removal of BrdU/BrdC incorporated DNA as previously described^73^.

### Telomere Fluorescence *in situ* Hybridization (Telo-FISH), imaging, quantifications

#### Protocol

Slides with metaphase chromosome spreads were prepared and hybridized with a fluorescently labeled telomere probe as previously described^65^. Briefly, slides were washed in 1x PBS for 5 min, dehydrated with an ice-cold ethanol series (75%, 85% and 100%) for 2 min each, air dried, and denatured in 70% formamide in 2x saline sodium citrate (SSC) at 75°C for 2 min, followed by a second ice-cold ethanol series, and air dried again. Probe hybridization mixture consisted of G-rich (TTAGGG-’3) peptide nucleic acid (PNA) telomere probe labeled with Cyanine-3 (Cy3; Biosynthesis) at a 5nM concentration in 36 μL of formamide, 12 μL of 0.5 M Tris-HCl, 2.5 μL of 0.1 M KCl, and 0.6 μl of 0.1 M MgCl2. Hybridization mixture was incubated at 75°C for 5 min and cooled on ice for 10 min, then 50 μL of mix was applied to each slide. Slides were coverslipped and hybridized at 37°C for 4 h. After hybridization, slides were washed five times at 43.5°C for three min each: washes one and two: 50% formamide in 2xSSC; washes three and four: 100% 2xSSC; and washes five and six: 2xSSC plus 0.1% Nonidet P-40. After washing, slides were counterstained with one drop of DAPI in Prolong Gold Antifade (ThermoFisher #P36931), coverslipped, and stored at 4°C for 24 h prior to imaging.

#### Image acquisition

Metaphase spreads (50 per patient/time point) were imaged at 100x mag on a Zeiss Axio Imager.Z2, Cool SNAP ES2 camera, and X-cite 120 LED lamp lightsource.

#### Individual telomere quantifications

Relative fluorescence intensity of individual telomeres was quantified using the ImageJ^74^ plugin Telometer (https://demarzolab.pathology.jhmi.edu/telometer/). Variation in Telo-FISH was controlled by assigning each patient a pair of slides made from BJ1 primary cells and BJ-hTERT cell lines. For each patient the slide preparation, Telo-FISH protocol, image acquisition and telomere quantifications were performed on the full time-course of samples and a pair of BJ1/BJ-hTERT controls (50 metaphases per control) at the same time and on the same respective days. Mean telomere length was quantified for each pair of control samples yielding a ratio for standardizing patients’ telomere values as previously described^75^.

### Telo-FISH data processing, feature engineering of short and long telomeres

#### Processing individual telomere length data

For each patient, outliers were removed from individual telomere length data per sample by omitting measurements three standard deviations from the mean. For samples with fewer individual telomere length measurements than the theoretical number (human cells, 50 metaphase spreads), missing telomere values were imputed by randomly sampling measurements from the observed distribution of individual telomeres; randomly sampled telomeres were added up to the theoretical number of telomeres per sample.

#### Feature engineering short and long telomeres

Individual telomeres from the pre-IMRT nonirradiated time point were split into quartiles, designating telomeres in the bottom 25% in yellow, the middle 50% in blue, and top 25% in red. Quartile cut-off values, established by the pre-IMRT non-irradiated sample’s distribution (values that separate quartiles), were applied to subsequent time points to feature engineer the relative shortest (yellow), mid-length (blue), and longest (red) individual telomeres per time point.

### Statistical and clustering analyses of Telo-FISH data

Statistical and clustering analyses were conducted with Python in Jupyter notebooks (see Code availability). With the statsmodels library^76^, mean telomere length and numbers of short and long telomeres were analyzed with a repeated measures ANOVA and post-hoc Tukey’s HSD test (two-tailed p values for both tests). Analyses were performed on all patients (n=14, less patient ID 13; 3 months post-IMRT sample failed to culture) and all four time course samples. A square root transformation was performed on numbers of short and long telomeres prior to statistical analysis. Ordinary least squares linear regression was performed with the scikit-learn LinearRegression tool. Hierarchical clustering analyses were performed on z-score normalized data using the scipy library with a single linkage method and Pearson correlation metric. Pearson correlations between patients’ longitudinal measurements of telomere length and complete blood count data was done with Python.

### XGBoost models with individual telomere length data, randomized hyperparameter search, cross validation

XGBoost models, model hyperparameter tuning, and cross validation tools were performed in Python through the scikit-learn API^77^. XGboost model features were individual telomere length values and sample labels denoted pre-IMRT sample origin (non-irradiated, *in vitro* irradiated), which were encoded as 0/1. Model hyperparameters were tuned using a randomized search with RandomizedSearchCV. For models predicting mean telomere length at late post-IMRT, final model hyperparameters were modified as follows: n_estimators=200, max_depth=7, learning_rate=0.2, objective =‘reg:squarederror’, random_state=1. For models predicting short and long telomeres at late post-IMRT, final model hyperparameters were similar as for mean telomere length, with max_depth=6. Five-fold cross validation was performed with cross_val_score and a negative mean absolute error metric.

### directional Genomic Hybridization (dGH), image acquisition, data processing

#### Protocol

High-resolution detection of chromosome aberrations (inversions, translocations) was performed with directional Genomic Hybridization (dGH) whole chromosome (Cy3) and subtelomere (Cy5) paints to chromosomes 1, 2, and 3 (KromaTiD Inc.) as previously described^57^. Briefly, slides were submersed in Hoechst 33258 (Millipore Sigma #B1155) for 15 min, photolyzed for 35 min using a SpectroLinker UV Crosslinker (365 nm UV), and treated with exonuclease III (New England Biolabs #M0206L) for 30 min. Paint hybridization mixture was applied to slides, which were then coverslipped, sealed with rubber cement, and denatured at 70°C for three min. Slides were hybridized for 24 h at 37°C, followed by five washes in 2xSSC at 43.5°C. After washing, slides were counterstained with one drop of DAPI in Prolong Gold Antifade (ThermoFisher #P36931), coverslipped, and stored at 4°C for 24 h prior to imaging.

#### Image acquisition

Metaphase spreads (30 per patient/time point) were imaged/scored at 63x mag on a Zeiss Axio Imager.Z2, Cool SNAP ES2 camera, and X-cite 120 LED lamp lightsource.

#### Data processing

Counts of chromosome aberrations were adjusted for clonality, where identical aberrations between cells for a patient’s given time point were noted but scored only once.

### Statistical and clustering analyses of chromosome aberrations (dGH)

Statistical and clustering analyses were conducted with Python in Jupyter notebooks (see Code availability). With the statsmodels library, average chromosome aberration frequencies were analyzed with a repeated measures ANOVA and post-hoc Tukey’s HSD test (two-tailed p values for both tests). Analyses were performed on all patients (n=14, less patient ID 13; 3 months post-IMRT sample failed to culture) and all time course samples (4). Ordinary least squares linear regression was performed with the scikit-learn LinearRegression tool. Hierarchical clustering analyses were performed on z-score normalized data using the scipy library with a single linkage method and Pearson correlation metric. Pearson correlations between patients’ longitudinal measurements of average chromosome aberration frequencies and complete blood count data was done with Python.

### XGBoost model design with chromosome aberrations

XGBoost models, model hyperparameter tuning, and cross validation were accessed in Python via the same manner as described for Telo-FISH data above. XGboost model features were counts of scored chromosome aberrations per cell, with sample labels denoting pre-IMRT sample origin (non-irradiated, *in vitro* irradiated; encoded as 0/1). Model hyperparameters were tuned using a randomized search with RandomizedSearchCV; models were ultimately non-performant. Final model hyperparameters (used with all chromosome aberrations) were: n_estimators=200, max_depth=15, learning_rate=0.1, objective=‘reg:squarederror’, random_state=0. Five-fold cross validation was performed with a negative mean absolute error metric.

## Notes

https://github.com/Jared-Luxton/radiation-therapy-machine-learning

