## Supplementary figures and tables for "Telomere length and chromosomal instability for predicting individual radiosensitivity and risk via machine learning"

### Supplementary figures:

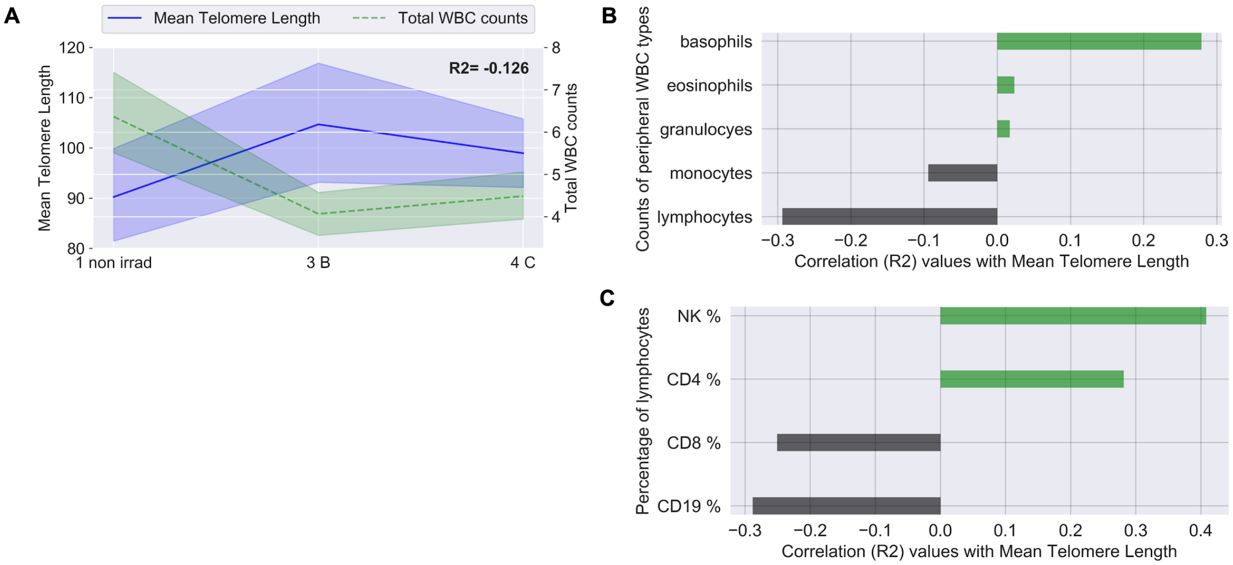


**SUPPLEMENTARY Fig 1. Correlations between telomere length, peripheral white blood cells, and lymphocytes.** Mean telomere length (Telo-FISH) plotted longitudinally against peripheral white blood cell (WBC) counts (thousands per microliter) from complete blood count tests for all patients; and longitudinal correlations between mean telomere length and counts of WBC types, and proportions of lymphocyte cell types. 1 non irrad: pre-IMRT non-irradiated; 3 B: immediate post-IMRT; 4 C: 3 months post-IMRT. Pearson correlation R2 values were calculated between longitudinal values, as shown bolded in (A), on a per patient basis. Correlations between mean telomere length and WBC counts **A**); center lines denote medians, lighter bands denote confidence intervals. Correlations between mean telomere length and five main WBC types **B**), and proportions of lymphocyte cell types **C**).

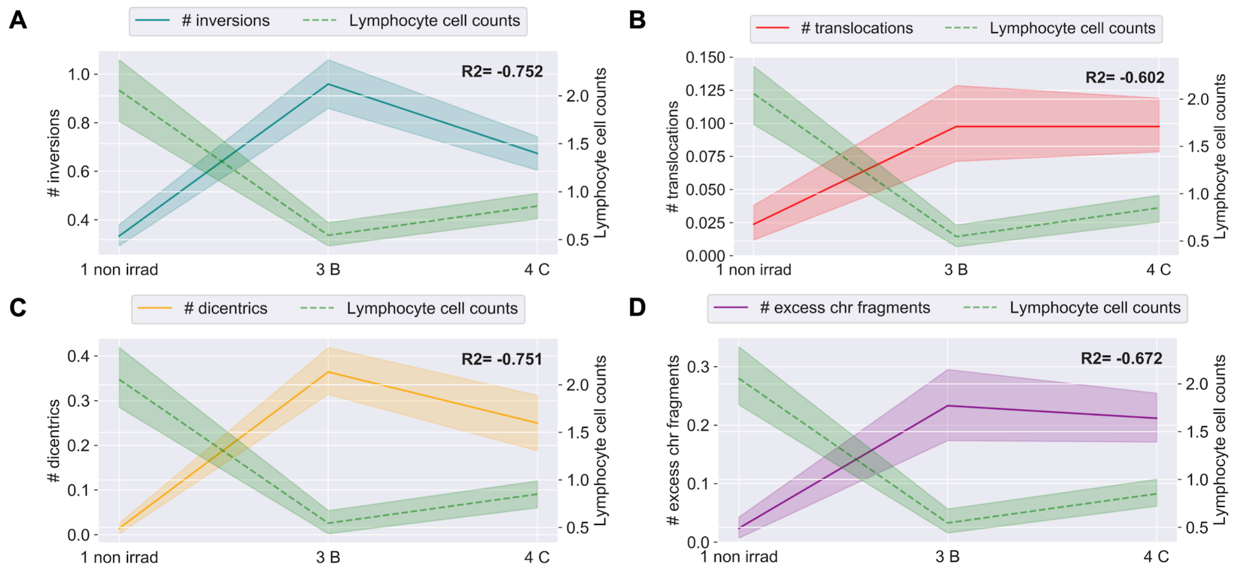


**SUPPLEMENTARY Fig 2. Correlations between chromosome aberrations and peripheral blood lymphocytes.** Average frequencies of chromosome aberrations plotted longitudinally against lymphocyte cell counts (thousands per microliter) from complete blood count tests for all patients. 1 non irrad: pre-IMRT non-irradiated; 3 B: immediate post-IMRT; 4 C: 3 months post-IMRT. Excess chr fragments: counts of chromosome fragments per cell after subtracting 1 count per n observed dicentrics. Center lines denote medians, lighter bands denote confidence intervals. Pearson correlation R2 values were calculated between plotted values on a per patient basis and noted in bold on each graph. **A**) Inversions, **B**) translocations, **C**) dicentrics, **D**) chromosome fragments and lymphocyte cell counts.

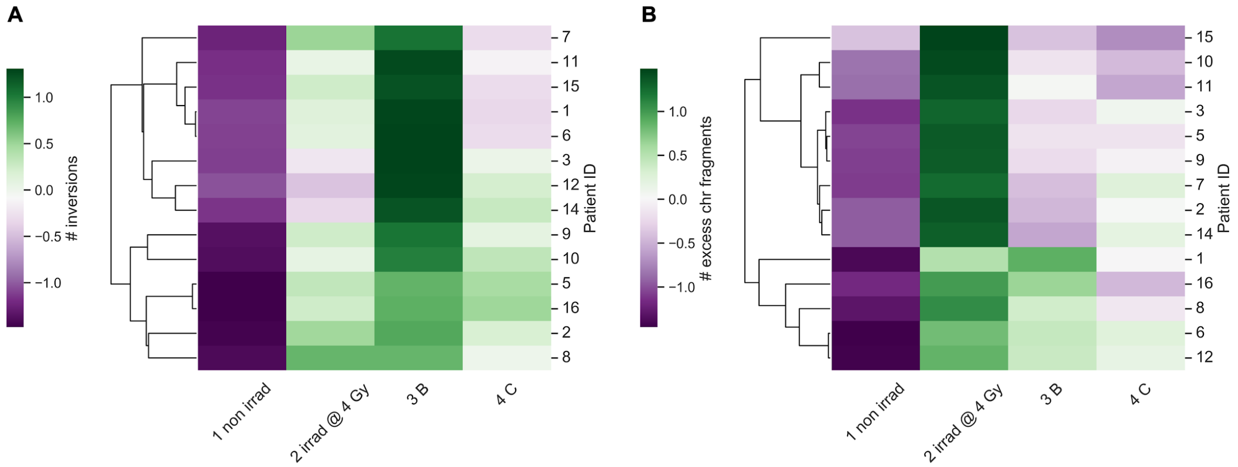


**SUPPLEMENTARY Fig 3. Clustering of patients by inversions and chromosome fragments (deletions).** Hierarchical clustering of patients by longitudinal changes in chromosome aberrations scored by directional Genomic Hybridization (dGH). 1 non irrad: pre-IMRT non-irradiated; 2 irrad @ 4 Gy: pre-IMRT *in vitro* irradiated; 3 B: immediate post-IMRT; 4 C: 3 months post-IMRT. Excess chr fragments: counts of chromosome fragments per cell after subtracting 1 count per n observed dicentrics. Patients were clustered by inversions **A**) and chromosome fragments **B**) (z-score normalized). Patient ID 13 not clustered; 3 months post-IMRT sample failed to culture.

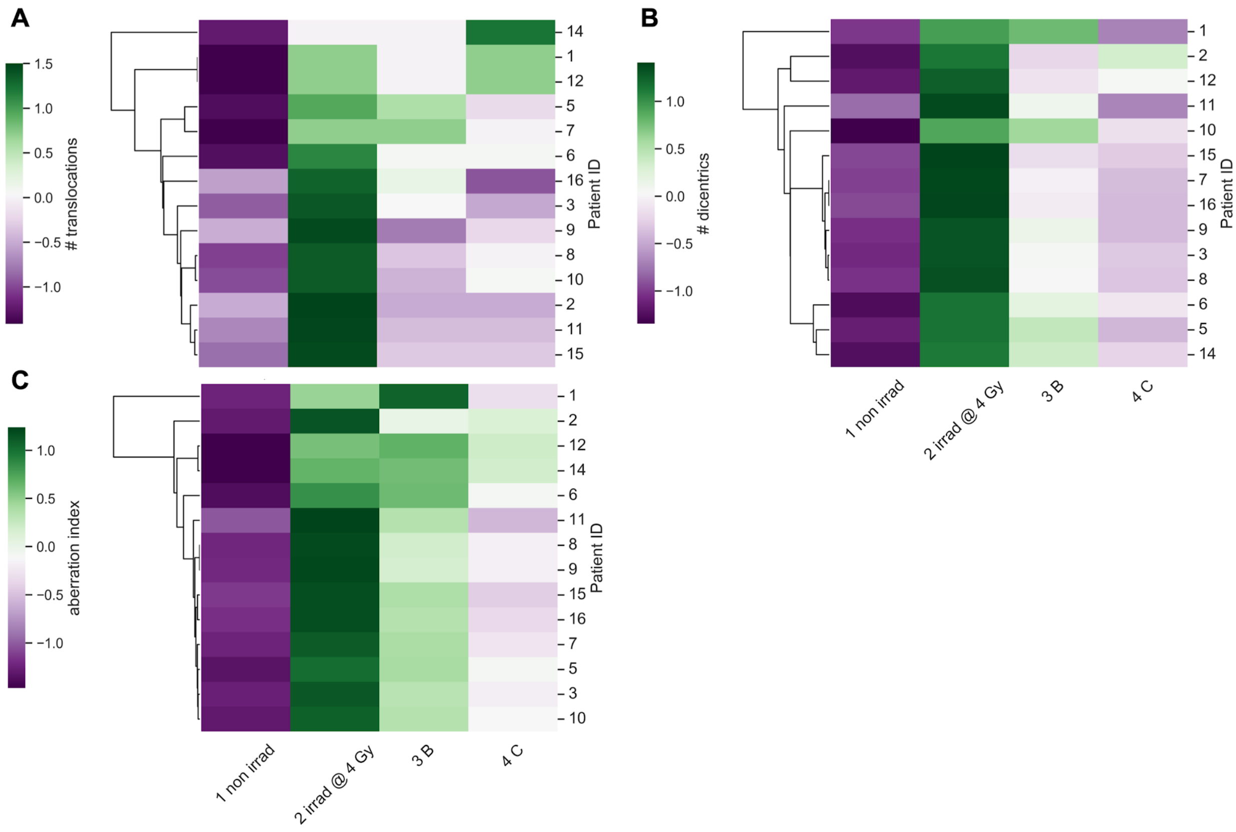


**SUPPLEMENTARY Fig 4. Chromosome aberrations generally failed to cluster patients.** Hierarchical clustering of patients by longitudinal changes in chromosome aberrations scored by directional Genomic Hybridization (dGH). 1 non irrad: pre-IMRT non-irradiated; 2 irrad @ 4 Gy: pre-IMRT *in vitro* irradiated; 3 B: immediate post-IMRT; 4 C: 3 months post-IMRT. Aberration index is created by summing all aberrations (inversions, translocations, dicentrics, chromosome fragments) per cell. Patients were clustered by translocations **A**), dicentrics **B**), and aberration index **C**) (z-score normalized). Patient ID 13 not clustered; 3 months post-IMRT sample failed to culture.

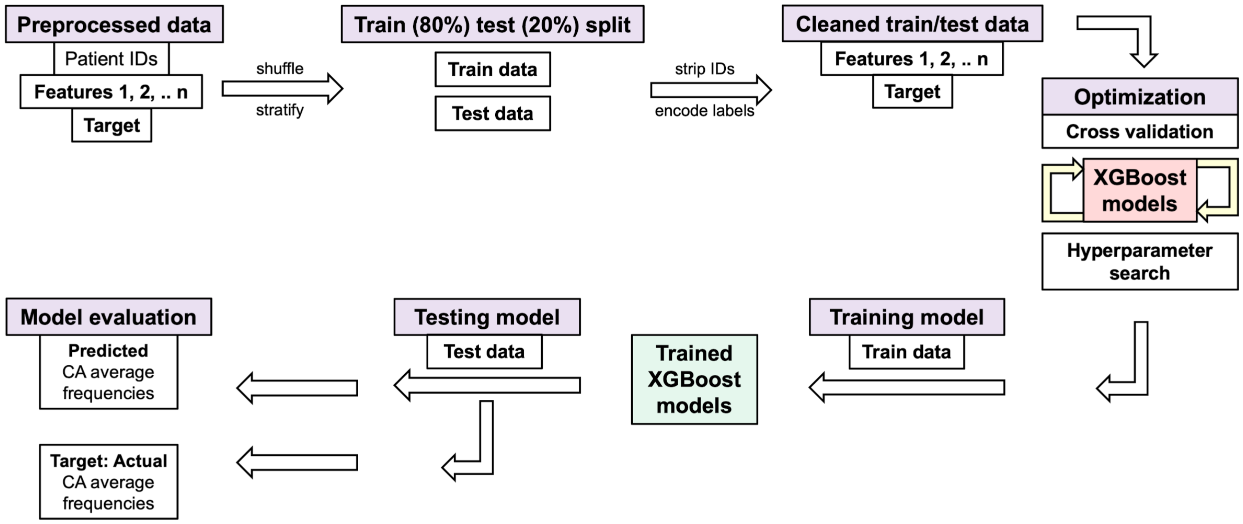


**SUPPLEMENTARY Fig 5. Processing of chromosome aberration data for XGBoost models.** Schematic for machine learning pipeline using chromosome aberration (CAs) data. Preprocessed data: Feature 1: pre-IMRT counts of scored CAs; Feature 2: pre-IMRT sample labels (non-irradiated, *in vitro* irradiated, encoded as 0/1); Feature n: represents pre-IMRT counts of multiple types of CAs (for aberration index). Target: Late post-IMRT average frequencies of CAs (either specific aberration type or aberration index). Data is randomly shuffled and stratified (by patient ID and pre-IMRT sample origin) and split into training (80%) and testing (20%) datasets; patient IDs are stripped after splitting. Five-fold cross validation was used, and models were evaluated with Mean Absolute Error (MAE) and $R^2$ between predicted and true values in the test set. See Materials and Methods and Code availability for model parameters and implementations in Python.

### Supplementary tables:

A

| patient id | pre-therapy sample origin | individual telomeres (RFI) | 4 C telo means |
|---|---|---|---|
| 1 | 1 non irrad | 52.79329603949808 | 99.34629891451401 |
| 1 | 2 irrad @ 4 Gy | 100.30726247504634 | 99.34629891451401 |
| 1 | 1 non irrad | 59.12849156423784 | 99.34629891451401 |
| 1 | 2 irrad @ 4 Gy | 106.64139157520613 | 99.34629891451401 |
| 1 | 1 non irrad | 69.68715077213746 | 99.34629891451401 |
| 1 | 2 irrad @ 4 Gy | 107.69724693733689 | 99.34629891451401 |

B

| encoded sample origin | individual telomeres (RFI) | 4 C telo means |
|---|---|---|
| 1.0 | 71.84704355757034 | 90.6803515449468 |
| 0.0 | 58.01948086775996 | 108.91532697997721 |
| 0.0 | 125.05216035895008 | 93.35225326745208 |
| 0.0 | 99.84003125432304 | 93.35225326745208 |
| 0.0 | 157.34096506511176 | 108.91532697997721 |
| 1.0 | 59.127900279322205 | 99.34629891451401 |

C

| patient id | pre-therapy sample origin | individual telomeres (RFI) | 4 C # short telos |
|---|---|---|---|
| 1 | 1 non irrad | 52.79329603949808 | 372 |
| 1 | 2 irrad @ 4 Gy | 100.30726247504634 | 372 |
| 1 | 1 non irrad | 59.12849156423784 | 372 |
| 1 | 2 irrad @ 4 Gy | 106.64139157520613 | 372 |
| 1 | 1 non irrad | 69.68715077213746 | 372 |
| 1 | 2 irrad @ 4 Gy | 107.69724693733689 | 372 |

D

| encoded sample origin | individual telomeres (RFI) | 4 C # short telos |
|---|---|---|
| 0.0 | 39.80714575487005 | 319.0 |
| 0.0 | 84.7669312523909 | 2028.0 |
| 0.0 | 48.569832356338225 | 372.0 |
| 1.0 | 99.34779587017763 | 829.0 |
| 1.0 | 104.85784735429183 | 319.0 |
| 1.0 | 92.25878757956735 | 124.0 |

E

| patient id | pre-therapy sample origin | individual telomeres (RFI) | 4 C # long telos |
|---|---|---|---|
| 1 | 1 non irrad | 52.79329603949808 | 1987 |
| 1 | 2 irrad @ 4 Gy | 100.30726247504634 | 1987 |
| 1 | 1 non irrad | 59.12849156423784 | 1987 |
| 1 | 2 irrad @ 4 Gy | 106.64139157520613 | 1987 |
| 1 | 1 non irrad | 69.68715077213746 | 1987 |
| 1 | 2 irrad @ 4 Gy | 107.69724693733689 | 1987 |

F

| encoded sample origin | individual telomeres (RFI) | 4 C # long telos |
|---|---|---|
| 0.0 | 56.551567152220926 | 2026.0 |
| 0.0 | 103.18180673387677 | 2026.0 |
| 0.0 | 69.58478047400733 | 365.0 |
| 0.0 | 56.18104859876975 | 1078.0 |
| 1.0 | 137.72889825629034 | 1002.0 |
| 1.0 | 84.46927366319693 | 1987.0 |

**SUPPLEMENTARY Table 1. Example views of individual telomere length data matrices used to train XGBoost models.** XGBoost models were trained on 103,040 individual telomere length measurements (one telomere per row) (Telo-FISH) from pre-IMRT non-irradiated (1 non irrad) and *in vitro* irradiated (2 irrad @ 4 Gy) samples to predict 3 months post-IMRT (4 C) telomeric outcomes. Matrices represent examples of pre- (A/C/E) and post-processed (B/D/F) training data. Patient IDs are stripped after data is shuffled and stratified. The 'encoded sample origin' column contains numerical encodings denoting individual telomeres' pre-IMRT sample of origin (0: non-irradiated, 1: *in vitro* irradiated). XGBoost models were trained to predict mean telomere length (**A**/**B**) and numbers of short (**C**/**D**) and long (**E**/**F**) telomeres at 3 months post-IMRT with data in the format as shown.

**A**

| Average MAE of CV folds | Std dev of MAE of CV folds | MAE predicted vs. test values | R2 predicted vs. test values | N samples training data |
|---|---|---|---|---|
| 11.4602 | 1.6502 | 13.4903 | -0.8393 | 100.0 |
| 10.6657 | 0.4454 | 10.3646 | -0.2049 | 500.0 |
| 8.0423 | 0.486 | 7.9009 | 0.1788 | 1000.0 |
| 6.7089 | 0.3895 | 6.0449 | 0.5126 | 2000.0 |
| 4.8488 | 0.2224 | 4.642 | 0.7094 | 4000.0 |
| 3.9282 | 0.0988 | 3.7677 | 0.8215 | 8000.0 |
| 3.6385 | 0.0447 | 3.5413 | 0.851 | 16000.0 |
| 3.3792 | 0.0626 | 3.3483 | 0.8755 | 32000.0 |
| 3.2944 | 0.051 | 3.2521 | 0.881 | 64000.0 |
| 3.233 | 0.052 | 3.2596 | 0.8817 | 103040.0 |

**B**

| Average MAE of CV folds | Std dev of MAE of CV folds | MAE predicted vs. test values | R2 predicted vs. test values | N samples training data |
|---|---|---|---|---|
| 705.0956 | 48.4789 | 680.9499 | -0.5887 | 100.0 |
| 573.3922 | 25.5422 | 521.0982 | -0.0162 | 500.0 |
| 440.9283 | 22.7264 | 425.5251 | 0.2572 | 1000.0 |
| 366.4338 | 19.0126 | 326.1635 | 0.5396 | 2000.0 |
| 315.2925 | 5.9607 | 292.0579 | 0.6593 | 4000.0 |
| 269.2991 | 6.6633 | 260.4209 | 0.7433 | 8000.0 |
| 257.6623 | 3.5097 | 247.6769 | 0.7747 | 16000.0 |
| 243.5729 | 4.1386 | 241.8505 | 0.7987 | 32000.0 |
| 233.7408 | 5.251 | 231.1663 | 0.803 | 64000.0 |
| 236.2825 | 2.0593 | 234.1744 | 0.8112 | 103040.0 |

**C**

| Average MAE of CV folds | Std dev of MAE of CV folds | MAE predicted vs. test values | R2 predicted vs. test values | N samples training data |
|---|---|---|---|---|
| 1056.6558 | 219.1554 | 953.2471 | -0.4405 | 100.0 |
| 763.7998 | 38.9092 | 727.2706 | 0.0447 | 500.0 |
| 629.6607 | 49.9928 | 627.9304 | 0.2945 | 1000.0 |
| 548.353 | 24.8756 | 481.5782 | 0.5641 | 2000.0 |
| 409.3232 | 4.8234 | 415.0895 | 0.674 | 4000.0 |
| 382.1325 | 11.974 | 376.8821 | 0.7505 | 8000.0 |
| 353.0249 | 6.234 | 348.5064 | 0.7981 | 16000.0 |
| 343.0401 | 4.5386 | 329.2967 | 0.8128 | 32000.0 |
| 331.0765 | 3.7999 | 331.8519 | 0.813 | 64000.0 |
| 330.3521 | 2.0857 | 335.931 | 0.8191 | 103040.0 |

**SUPPLEMENTARY Table 2. Metrics of XGBoost models for predicting post-IMRT telomeric outcomes.** XGBoost models were trained on pre-IMRT individual telomere length measurements (Telo-FISH) to predict 3 months post-IMRT telomeric outcomes. Metrics assess model performance during (five) cross-fold validation (CV) (columns 1-2 from left) and when challenged with the test set (test) (columns 3-4 from left). Model performance was evaluated with mean absolute error (MAE) (std dev: standard deviation) across a range of samples in the training data (n=100 to 103,040). $R^2$: correlation metric. Metrics of XGBoost models for predicting 3 months post-IMRT (4 C) mean telomere length **A**), numbers of short **B**) and long **C**) telomeres.

**A**

| patient id | pre-therapy sample origin | # inversions | 4 C # inversions |
|---|---|---|---|
| 5 | 1 non irrad | 0 | 0.7083333333333334 |
| 11 | 2 irrad @ 4 Gy | 2 | 0.4583333333333333 |
| 1 | 1 non irrad | 0 | 0.5 |
| 9 | 1 non irrad | 0 | 0.7083333333333334 |
| 11 | 1 non irrad | 0 | 0.4583333333333333 |
| 16 | 1 non irrad | 0 | 0.7916666666666666 |

**B**

| encoded sample origin | # inversions | 4 C # inversions |
|---|---|---|
| 0.0 | 0.0 | 0.7083333333333334 |
| 1.0 | 0.0 | 0.5 |
| 1.0 | 0.0 | 0.5 |
| 0.0 | 1.0 | 0.7083333333333334 |
| 1.0 | 0.0 | 0.5 |
| 1.0 | 1.0 | 0.7916666666666666 |

**C**

| patient id | pre-therapy sample origin | # inversions | # translocations | # dicentrics | # excess chr fragments | 4 C aberration index |
|---|---|---|---|---|---|---|
| 9 | 1 non irrad | 0 | 0 | 0 | 0 | 1.125 |
| 7 | 2 irrad @ 4 Gy | 1 | 0 | 1 | 1 | 0.8333333333333334 |
| 11 | 2 irrad @ 4 Gy | 0 | 1 | 0 | 0 | 0.6666666666666666 |
| 1 | 1 non irrad | 0 | 0 | 0 | 0 | 0.9583333333333334 |
| 16 | 1 non irrad | 0 | 0 | 0 | 0 | 1.0833333333333333 |
| 6 | 1 non irrad | 0 | 0 | 0 | 0 | 1.375 |

**D**

| encoded sample origin | # inversions | # translocations | # dicentrics | # excess chr fragments | 4 C aberration index |
|---|---|---|---|---|---|
| 0.0 | 0.0 | 0.0 | 0.0 | 0.0 | 1.0833333333333333 |
| 0.0 | 0.0 | 0.0 | 0.0 | 0.0 | 0.9583333333333334 |
| 1.0 | 0.0 | 0.0 | 0.0 | 0.0 | 1.4166666666666667 |
| 0.0 | 0.0 | 0.0 | 0.0 | 0.0 | 1.2083333333333333 |
| 1.0 | 2.0 | 0.0 | 0.0 | 0.0 | 1.2083333333333333 |
| 0.0 | 0.0 | 0.0 | 0.0 | 0.0 | 0.9166666666666666 |

**SUPPLEMENTARY Table 3. Example views of chromosome aberration data matrices used to train XGBoost models.** XGBoost models were trained on chromosome aberration count data (one cell per row, n=672) from pre-IMRT non-irradiated (1 non irrad) and *in vitro* irradiated (2 irrad @ 4 Gy) samples to predict 3 months post-IMRT (4 C) chromosome aberration frequencies. Matrices represent pre- (A/C) and post-processed (B/D) training data. Patient IDs are stripped after data is shuffled and stratified. The 'encoded sample origin' column contains numerical encodings denoting cells' pre-IMRT sample of origin (0: non-irradiated, 1: *in vitro* irradiated). XGBoost models shown were trained to predict average inversion frequencies (**A**/**B**) and aberration index frequencies (**C**/**D**).

**A**

| Features | Target | Average MAE of CV folds | Std dev of MAE of CV folds | MAE predicted vs. test values | R2 predicted vs. test values |
|---|---|---|---|---|---|
| # inversions, encoded samples | 4 C # inversions | 0.1746 | 0.0445 | 0.2724 | -0.213 |
| # translocations, encoded samples | 4 C # translocations | 0.0412 | 0.0188 | 0.1327 | -0.3905 |
| # dicentrics, encoded samples | 4 C # dicentrics | 0.1171 | 0.0461 | 0.2508 | 0.0019 |
| # excess chr fragments, encoded samples | 4 C # excess chr fragments | 0.0939 | 0.0334 | 0.1787 | -0.1228 |
| all aberrations, encoded samples | 4 C aberration index | 0.2541 | 0.0496 | 0.5137 | -0.05 |

**B**

| Features | Target | Average MAE of CV folds | Std dev of MAE of CV folds | MAE predicted vs. test values | R2 predicted vs. test values |
|---|---|---|---|---|---|
| # inversions, encoded samples | 4 C # inversions | 0.1759 | 0.0332 | 0.2187 | -1.5965 |
| # translocations, encoded samples | 4 C # translocations | 0.0375 | 0.0096 | 0.1096 | -0.004 |
| # dicentrics, encoded samples | 4 C # dicentrics | 0.1167 | 0.0463 | 0.2023 | -0.0215 |
| # excess chr fragments, encoded samples | 4 C # excess chr fragments | 0.084 | 0.0103 | 0.202 | -0.1071 |
| all aberrations, encoded samples | 4 C aberration index | 0.3294 | 0.1258 | 0.3596 | -0.0256 |

**C**

| Features | Target | Average MAE of CV folds | Std dev of MAE of CV folds | MAE predicted vs. test values | R2 predicted vs. test values |
|---|---|---|---|---|---|
| # inversions, encoded samples | 4 C # inversions | 0.15 | 0.0573 | 0.1977 | -0.0709 |
| # translocations, encoded samples | 4 C # translocations | 0.0448 | 0.0145 | 0.0977 | -0.0346 |
| # dicentrics, encoded samples | 4 C # dicentrics | 0.1123 | 0.034 | 0.2177 | -0.0558 |
| # excess chr fragments, encoded samples | 4 C # excess chr fragments | 0.0681 | 0.0155 | 0.194 | -0.0372 |
| all aberrations, encoded samples | 4 C aberration index | 0.3255 | 0.0528 | 0.5078 | -0.0259 |

**SUPPLEMENTARY Table 4. Metrics of trained XGBoost models for predicting post-IMRT average frequencies of chromosome aberrations.** Multiple iterations of XGBoost models (A-C) were trained on pre-IMRT chromosome aberration counts per cell (n=672 cells) to predict late post-IMRT average chromosome aberration frequencies. Time points for pre-IMRT data were encoded (0/1: non-irradiated, *in vitro* irradiated). Metrics assess model performance during (five) cross-fold validation (CV) and when challenged with the test set (test). Model performance was evaluated with mean absolute error (MAE) (std dev: standard deviation). $R^2$: correlation metric. Performance of models with identical initializations and hyperparameters for predicting average frequencies of inversions , translocations, dicentrics, chromosome fragments, and aberration index are shown (**A**-**C**).
